# Comprehensive characterization of the viscoelastic properties of Bovine Submaxillary Mucin (BSM) hydrogels and the effect of additives

**DOI:** 10.1101/2024.01.17.576019

**Authors:** Hanna Rulff, Robert F. Schmidt, Ling-Fang Wei, Kerstin Fentker, Yannic Kerkhoff, Philipp Mertins, Marcus A. Mall, Daniel Lauster, Michael Gradzielski

## Abstract

This study presents a comprehensive characterization of the viscoelastic and structural properties of Bovine Submaxillary Mucin (BSM), which is widely used as a commercial source to conduct mucus-related research. We conducted concentration studies of BSM and examined the effects of various additives - NaCl, CaCl_2_, lysozyme, and DNA - on its rheological behavior. A notable connection between BSM concentration and viscoelastic properties was observed, particularly under varying ionic conditions. The rheological spectra could be well-described by a fractional Kelvin-Voigt Model with a minimum of model parameters. A detailed proteomics analysis provided insight into the molecular interactions within BSM, showing MUC19 as main component. Cryo-scanning electron microscopy allowed to visualize the network structure in relation to the rheological data. By elucidating the complex interplay between mucin concentration, environmental conditions, and viscoelastic properties, this research significantly contributes to the field of mucus research and lays an important basis for its further advancement.

## INTRODUCTION

Mucus is an essential component for mechanical and biological protection of many organisms, which covers epithelial surfaces. In human, mucus acts as a protective barrier for mucosal surfaces ^1^. Mucus is a complex hydrogel constituted by more than 95 % of water and (glyco)proteins, lipids, DNA, and salts ^2, 3^. Mucus owes its diverse and proper functioning to the interaction of its components, the most important being mucin glycoproteins. One of the central mucus features is its hydrogel-like viscoelastic characteristic, which is essential to limit diffusion of pathogenic particles, and to allow efficient transport via cilia-supported transport (mucociliary clearance) in the airways ^4^ or the peristaltic transport in the intestine. The hydrogel-gel like characteristic of mucus is shaped by mucins ^3, 5, 6^. Mucins are high molecular weight macromolecules showing extensive glycosylation, which allows high water-binding capacity, and together with the mainly disordered protein backbone structure hydrogel characteristics ^7–9^. Mucins form three-dimensional networks with a mesh-like structure, being responsible for the biophysical properties of mucus ^10–12^.

Native human mucus is a complex and heterogeneous biomaterial, which makes the generation of reproducible data challenging due to its natural polymeric dispersity within and among individual hosts ^3^ as well as the limited availability of mucus from human sources. Therefore, mucus from other sources possessing defined and reproducible properties and being available in larger quantities may provide suitable models for mucus research ^10, 13^. Here, the viscoelastic properties are a common feature of different mucus types arising from varying sources ^13, 14^. One commercially available mucus system is bovine submaxillary mucin (BSM), a native mucin, that is obtained by purification processes from extracts of fresh cattle submaxillary glands ^15^. However, BSM is not completely pure, and contains beside large quantities of glycoproteins also non-mucin proteins (for example bovine serum albumin (BSA)), DNA, lipids, and inorganic salts ^16^. BSM contains mucins ^17^ which are similar to human respiratory mucins ^18, 19^. They are known to confer viscoelastic properties to mucus and to be highly entangled polymers ^20^ that are dominated by repeating structural motifs that react sensitively to changes in pH and concentration of Ca^2+^ ions^21^. Accordingly, BSM can be considered as a model system for human mucus, having the advantage of being available in larger amounts of reproducible quality.

Such model systems are essential for gaining a fundamental understanding of the molecular origins of the viscoelastic properties of mucus and how they might be affected by changes of the solvent conditions (e. g. ionic strength, salt, etc.) or the presence of other components such as proteins or DNA. This knowledge of the mechanical response of mucus are crucial for gaining a deepened understanding, for instance of muco-obstructive lung diseases such as cystic fibrosis (CF) ^6, 22, 23^. Due to altered viscous and elastic properties of sputum in different stages of disease and therapy, measuring the rheological properties turned out to be a potential biomarker ^14, 24–26^. Noting this promising connection, BSM has already been studied to some extent. Thornton et al. designed technologies for the detection and quantification of mucins, also for BSM ^18^,while other groups gave insights in the BSM genome ^19, 27^. Lee et al. studied the behaviour of BSM on hydrophobic surfaces observing the formation of elastic films ^28^, also by changing the experiments environmental conditions ^29^. BSM was also tested to produce mucin-based hydrogels showing physiologically relevant properties by crosslinking it with different crosslinking reagents ^13, 14^. Furthermore, the commercially available mucins like porcine gastric mucin (PGM) and BSM were subjected to a characterisation and comparison with human sputum by Lee et al. They recommend using BSM as the currently most suitable commercially available mucin source for replicating saliva based on its surface adsorption and lubrication properties ^30^. Feiler et al. studied the viscoelastic properties of BSM and BSA upon adsorption to surfaces and their potential for applications in biomaterial coating, showing the ability of BSM to form viscoelastic and more rigid layers ^31^. In a mixture of BSM with β-lactoglobulin, BSM was found to decrease the viscoelastic moduli, and thus the stability of the viscoelastic network studied here at an air-liquid interface ^32^. A study from Seller et al. gave insights in the gel-formation of pig and sheep submaxillary mucins (PSM and SSM) influenced by their intact polymeric structure ^33^. The rheological behavior of commercially available PGM was subjected to a comparison to isolated natural porcine gastric mucus, indicating a more robust network formed by the natural mucus ^34^. Additionally, the influence of e.g. ionic strength, pH, and concentration on the rheological properties of commercially available PGM as well as isolated porcine gastric and duodenal mucins were studied, showcasing its adaptable mechanical properties in different environments ^35, 36^.

Ribbeck et al. provided valuable insights into the complex and adaptable rheological properties of purified MUC5AC from fresh pig stomach scrapings, highlighting the influence of environmental factors such as pH, salt, and surfactants on their viscoelastic behavior ^37^.

Despite comprehensive work on the rheological properties of mucins, this important class of hydrogels is far from being understood, which motivated us to provide a thorough characterization of the physico-chemical properties of BSM as a biological model material, with a particular focus on its viscoelastic properties. BSM can provide a potential model system for e.g. respiratory mucus, offering valuable insights in transport study applications. We performed a comprehensive investigation of the rheological properties of BSM as a function of concentration, and studied effects arising from the presence of salts, lysozyme as a model protein, and DNA. A detailed molecular characterization of BSM was performed, mainly by oscillatory and shear flow rheology experiments conducted on corresponding series of samples, from which systematic correlations between the viscoelastic properties and the composition of the samples could be deduced. The interpretation of the rheological data was done by employing a fractional Kelvin-Voigt model, which delivers a fitting description of the observed experimental data with a small number of parameters and thereby constitutes a well-adapted theoretical model to describe the rheological behavior of mucins.

## MATERIALS AND METHODS

### Preparation of mucus from bovine submaxillary mucin

BSM (Mucin, Bovine Submaxillary Gland, Merck KGaA, Darmstadt, Germany, LOT 3776068, and LOT 3829388 was dissolved in Dulbecco’s phosphate buffered saline (DPBS) without calcium and magnesium (containing 8 mg/mL sodium) (Lot No. RNBJ1061, Sigma-Aldrich) for 45 minutes at room temperature with a magnetic stirrer at the lowest speed (100 rotations per minute) to prevent the formation of bubbles. We prepared different concentrated BSM solutions with 20, 40, 60, 100, 140 mg/mL (= 2.0, 4.0, 6.0, 10.0, 14.0 % w/v). pH measurements were performed directly after preparation using a micro-electrode (InLab Micro, Mettler Toledo). After this the rheology of the samples was measured.

### Measurement of the calcium and sodium content in BSM with Inductively Coupled Plasma - Optical Emission Spectrometry (ICP-OES)

For the BSM solutions of both batches (LOT 3776068 and LOT 3829388) the content of calcium and sodium was quantified by ICP-OES using a Varian ICP-OES 715 ES spectrometer. Details of the procedure are given in SI.

### Detection of the DNA content in BSM solution

The DNA content determination protocol was adapted from published literature ^38^, and the DNA content was determined based on the reaction of 3,5-diaminobenzoic acid dihydrochloride (DABA; TCI) and aldehyde group in DNA. The DNA reaction solution comprised 20% w/v DABA in milli Q water. 30 μL of BSM samples (LOT: 3678870) was mixed with 30 μL DNA detection solution and followed with incubation at 60 °C for 1 hour. The reaction was quenched by the addition of 1 ml of 1.75 M HCl. Fluorescence intensity was measured with Tecan M200 pro at excitation and emission wavelengths of 413 and 512 nm, respectively. The DNA concentration in BSM samples was calculated from the calibration curve which was generated with using known concentrations of DNA solution from salmon testes (Sigma-Aldrich).

### Measurement of protein concentration in BSM solution

The protein concentration determination procedure was followed with Pierce BCA protein assay kit manual (Thermo Fisher). 25 μL of BSM samples in transparent 96-well plate was mixed with 200 μL working reagent which is composed with 1.96% v/v of reagent B and 98.04% v/v reagent A. After incubation at 37 °C for 30 minutes, absorbance intensity was measured with Tecan M200 pro at wavelengths of 562 nm. The protein concentration in BSM samples was calculated from the calibration curve which was generated using known concentrations of bovine serum albumin (BSA) solution (Thermo Fisher). The protein concentration was 85.55 mg/mL in a 100 mg/mL BSA solution.

### Addition of salts, DNA and lysozyme to a 100 mg/mL BSM solution

To study the effect of different additives on the viscoelastic properties of BSM (LOT: 3829388) we prepared 100 mg/mL BSM solutions using the previously described method and added the following substances: Six different concentrations of calcium chloride (as calcium chloride dihydrate, Merck KGaA, ≥99.0 %, C7902) 2.7, 6.0, 11.0, 21.0, 51.0, and 75.0 mM in a 100 mg/mL BSM solution, which was prepared with 20 mM HEPES-buffer (2-(4-(2-Hydroxyethyl)-1- piperazinyl)-ethansulfonsäure) (Sigma-Aldrich, ≥99.5 %, H3375) as calcium chloride is not soluble in DPBS-buffer. Four different concentrations of sodium chloride (Sigma-Aldrich, ≥99.5 %, S9888) 0.2, 0.4, 2.2, and 5.2 M in a 100 mg/mL BSM solution; see SI for detailed procedure. Four different concentrations of desoxyribonucleic acid (DNA) single stranded from salmon testes (D9156, Sigma-Aldrich) 5 mg/mL, 10 mg/mL, 15 mg/mL and 20 mg/mL were present in a 100 mg/mL BSM solution. Lysozyme (Muramidase from hen egg white, Roche Diagnostics GmbH, Mannheim, ≥95.0 %) was added in concentrations of 1.0, 2.0, and 10.0 mg/mL in a 100 mg/mL BSM solution. All solutions were well stirred for two hours and stored in the fridge for 24 hours. Afterwards, they were let adjust to room temperature, the pH-values were measured as stated above, and are given in each results section. Then the rheological measurements were performed.

### Rheology

All rheology measurements were performed on an Anton-Paar MCR 502 WESP temperature-controlled rheometer using a cone-plate geometry with a cone angle of 1 ° and a cone diameter of 25 mm. The gap was set at 48 µm. The sample was transferred onto the lower static plate of the rheometer with a non-electrostatic spatula and the upper cone was lowered slowly. The measurements were performed at 25 °C under saturated atmosphere using a solvent trap whose reservoir was filled with 2 mL MilliQ water, to avoid evaporation effects. After reaching a steady temperature of 25 °C and waiting for five minutes measurements were started with the amplitude sweep.

The linear viscoelastic (LVE) region and the critical deformation were determined by amplitude sweeps performed at a constant frequency of 1 Hz and covering a range of strain amplitudes between 0.01 % and 100 %. The LVE region represents the range of amplitude strain values over which the viscoelastic parameters are independent of the applied forces ^39^. Based on the amplitude sweeps (see Figure S3) the amplitude of the oscillatory experiments was set to 1 % strain and the frequency was varied between 0.05 Hz and 50 Hz. The frequency sweep was first performed from low to high and thereafter from high to low frequencies and they were interpreted regarding the viscoelastic material properties in terms of the storage (G’) and the loss (G’’) modulus, or alternatively via the phase angle δ (°) and the complex viscosity η* (Pa·s), which are directly related to each other via:

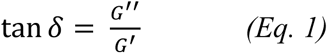

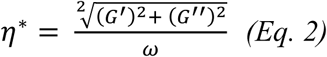

A fully elastic material is showing a phase angle of δ = 0°, while for fully viscous behavior a phase angle δ = 90° is observed. According to eq. 1 for δ < 45° the elastic properties dominate, while for predominant viscous behavior one has δ > 45°. The complex viscosity is another parameter to formulate the viscoelastic properties of a material, which can be compared directly to the normal shear viscosity, which was also studied in constant shear experiments, in which we varied the shear rate from 0.1 1/s to 1000 1/s, going from low to high and afterwards from high to low shear rates, in order to check for potential hysteresis effects. Data analysis was performed with GraphPad Prism version 10.1.2 (GraphPad Software, San Diego, CA; United States) and Origin, version 2021b (OriginLab Corporation, Northampton, MA, USA).

### Fitting of Rheological Data

Fitting of the frequency-dependent moduli *G*’ and *G*’’ was performed in Python using the lmfit package, which employs a non-linear least-squares optimization routine ^40^. Model functions were written to return a one-dimensional array containing *G*’ and *G*’’, meaning both moduli are fitted at the same time. The minimized quantity is defined as

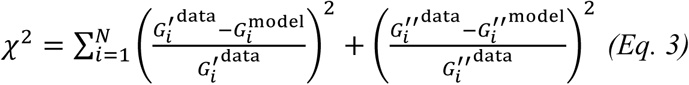

to ensure all data points are considered equally. At high frequencies there can be artefacts caused by sample and instrument inertia ^41^. For this reason, only angular frequencies *ω* < 50 rad/s = 7.96 Hz were considered for the fit.

### Cryo-Scanning Electron Microscopy (cryo-SEM)

A small drop of a 100 mg/mL BSM solution of each batch was plunged into nitrogen slush at atmospheric pressure, freeze-fractured at -180 °C, etched for 60 s at -98 °C and sputtered with platinum in a Gatan Alto 2500 cryo-preparation chamber. After transfer into the S-4800 cryo-SEM (Hitachi, Tokyo, Japan) the morphology of the BSM was evaluated at an acceleration voltage of 2.0 kV.

For the quantitative SEM evaluation ten images of each batch of BSM solution were included in the analysis. The included images were taken at magnifications of 20.000 x, 40.000 x, 80.000 x and 100.000 x. The SEM images were smoothed by applying a gaussian filter with a sigma radius of 1 pixel. Images were automatically binarized by an Otsu threshold ^42^ to separate them into the fiber network (bright) and pores (dark). The pores were analyzed with Fiji’s Particle Analyzer ^43^. Holes on the edges of the images were excluded from analysis to avoid artifacts. The maximal and minimal Feret diameter ^44–46^ of each pore was automatically quantified and manually verified. The distribution of minimal Feret diameters of all pores per sample was visualized in a violin plot with the mean value and the single standard deviation indicated as lines within the violin plot. Details of the procedure are given in SI (Figure S1).

### Proteomic measurements

BSM (LOT: 3829388) was dissolved in PBS at a concentration of 4 mg/mL. 3 aliquots à 100 µg mucine were reduced and alkylated in SDS buffer (2% SDS, 50 mM Tris-HCl pH 8, 0.5 mM EDTA, 75 mM NaCl, 10 mM DTT, 40 mM CAA final concentration) by incubating for 10 min at 95 °C. After cooling down bezonase (10.25 U, Merck) was added and samples incubated for 15 min at room temperature. Subsequently, single-pot, solid-phase-enhanced sample preparation (SP3 clean-up) was performed ^47^. N-glycosylation was removed by a 1-hour incubation with PNGase F (500 U, NEB) and following digest with sequence-grade trypsin (Promega) and lysyl endopeptidase (LysC, Wako) at a 1:50 enzyme:substrate ratio overnight at 37 °C. The digest was stopped by adding trifluoroacetic acid to a final concentration of 1%. The supernatant was collected, and peptides desalted using C18 stage tips ^48^. Peptides were reconstituted in 3% acetonitrile with 0.1% formic acid. 0.4 µg of peptides were separated on an EASY-nLC 1200 System on a reversed-phase column (20 cm fritless silica microcolumns with an inner diameter of 75 µm, in-house packed with ReproSil-Pur C18-AQ 1.9 µm resin (Dr. Maisch GmbH)) using a 98 min gradient with a 250 nl/min flow rate of increasing Buffer B (90% ACN, 0.1% FA) concentration (from 2% to 60%).

Separated peptides were analyzed on an Orbitrap Q Exactive HF-X mass spectrometer (Thermo Fisher Scientific) running on data dependent acquisition (DDA) mode with a 60 k MS1 resolution, AGC target of 3 × 10^6^ ions, and a maximum injection time of 10 ms, choosing the top 20 ions for MS2 scans with 15 K resolution, AGC target of 1 × 10^5^ ions and maximum injection time of 22 ms.

Database search was performed using MaxQuant (V 2.0.3.0) using Uniprot database for bovine proteins (downloaded 2022-09). Oxidation (M), N-terminal acetylation, and deamidation (N, Q) were set as variable modification and carbamidomethyl (C) as fixed modification. Match between runs, label-free quantification, and iBAQ algorithms were applied. Data are available via ProteomeXchange with identifier PXD048381.

Downstream analysis was done in R (V 4.2.2). Proteins were filtered for “reverse” and “only-identified by site”. Proteins identified by the contaminants list were removed if they did not originate from current bovine or cattle databases. Only proteins which were quantified in all of the replicates were considered for further analysis.

## RESULTS AND DISCUSSION

### Comparison of two different batches of pure BSM solution

First, we wanted to determine the dependence of data obtained from rheological characterization studies on the choice of BSM batch, as a batch-to-batch variability was an issue noted in the literature for commercially available mucus systems ^19, 27, 30, 49, 50^. Therefore, we used different batches of BSM, provided by Merck KgaA, and prepared by the method of Nisizawa K. and Pigman W. ^15^ and indicated by different LOT numbers.

From each batch 100 mg/mL BSM solutions in DPBS were prepared and underwent oscillatory rheological shear measurement. From both BSM batches the loss modulus G’’ dominated the storage modulus G’ within the investigated frequency range (Figure 1). Both batches showed very similar behavior in their frequency dependent trend of G’ and G’’ by increasing slowly with increasing frequency (Figure 1A). Notably, at lower frequencies G’ becomes relatively more prominent, as seen from the ratio G’/G’’, a comportment that is rather uncommon for viscoelastic systems (Figure 1B). This type of rheological behavior clearly indicated gel-like properties in the investigated frequency range but with a distinct viscous component. However, as seen by the rather low values of the moduli in the range of 0.5 to 20 Pa these were rather soft gel properties. The absolute values differed systematically by about 10 – 50 % (Figure 1B), which means that the selection of a given batch did not largely change the behavior, but certainly had a non-negligible effect on the absolute viscoelastic values. However, when looking at the ratio G’/G’’ (Figure 1B) the differences became very small, which means that the absolute values of G’ and G’’ depend on the batch, but the relative viscoelastic properties to a much lesser extent. Both samples did not differ in their pH, both having values of 7.4.

**Figure 1.**
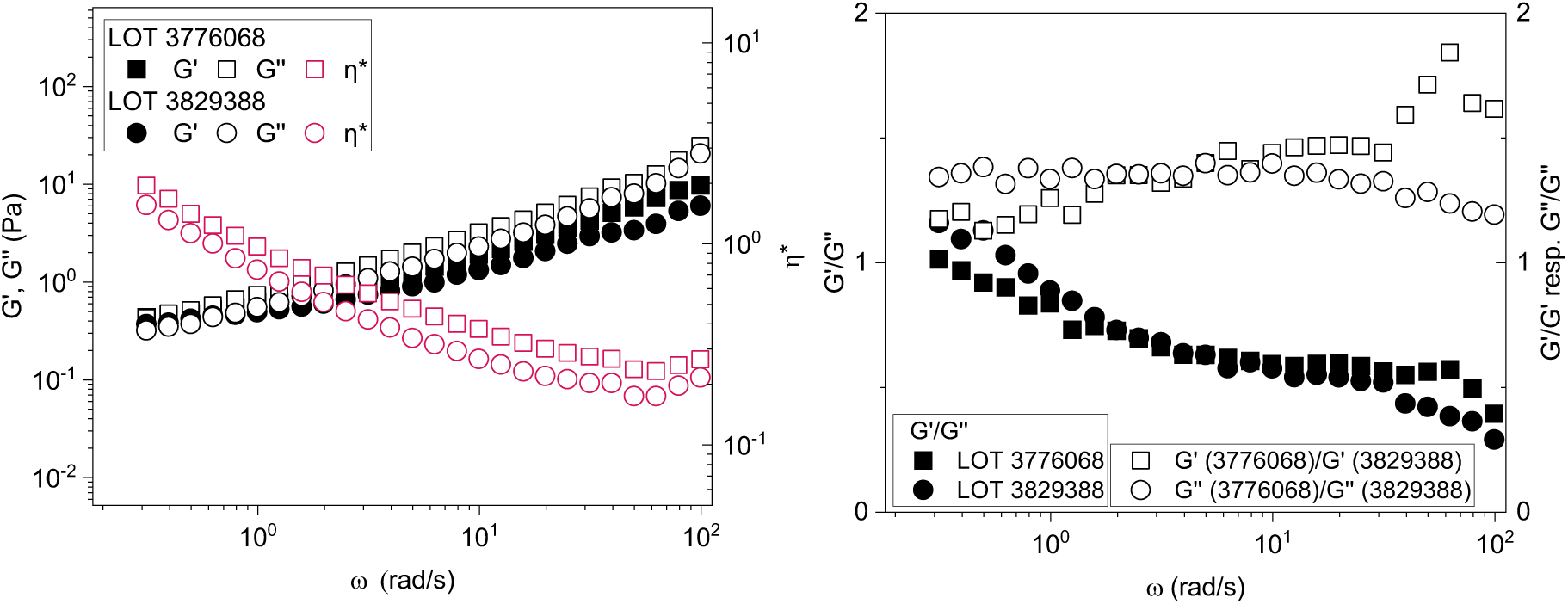
Viscoelastic properties of two different batches of BSM. Storage modulus G’ and loss modulus G’’ (Pa) as well as complex viscosity η* of 100 mg/mL BSM solutions were measured as function of angular frequency (rad/s) at 25 °C (data are shown as mean values of n=3 measurements) (A). Ratio of G‘/G‘‘ for the given two batches, as well as ratio of G’/G’ and G’’/G’’ for the two different batches as function of angular frequency (rad/s) (B).

Overall, the differences of the viscoelastic properties between the two investigated BSM batches were negligible small, as were their mesh sizes (shown as min Feret diameter [nm] in Figure 2C) obtained from a structured analysis of the cryo-SEM images.

**Figure 2A.**
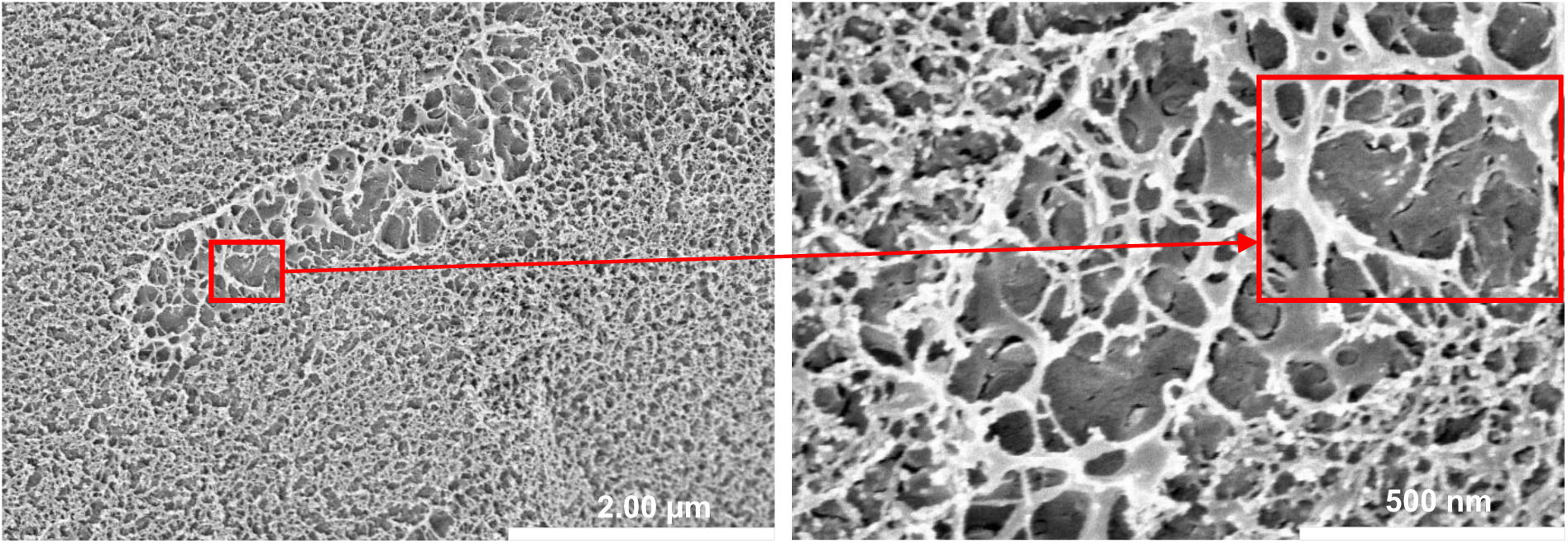
Exemplary SEM images. Cryogenic-scanning electron micrographs (cryo-SEM) at 20.000 (left) × and 80.000 (right) × magnification of a 100 mg/mL BSM solution, Batch-Nr. LOT 3776068.

**Figure 2B.**
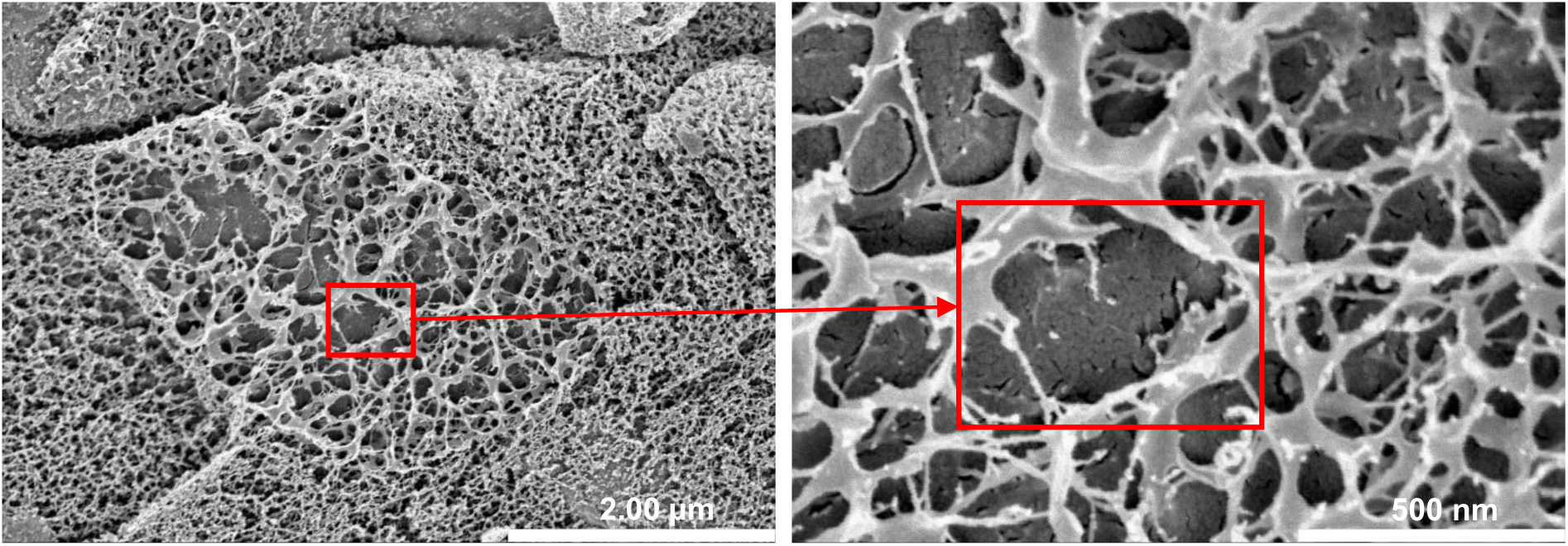
Exemplary SEM images. Cryogenic-scanning electron micrographs (cryo-SEM) at 20.000 (left) × and 80.000 (right) × magnification of a 100 mg/mL BSM solution, Batch-Nr. LOT 3829388.

**Figure 2C.**
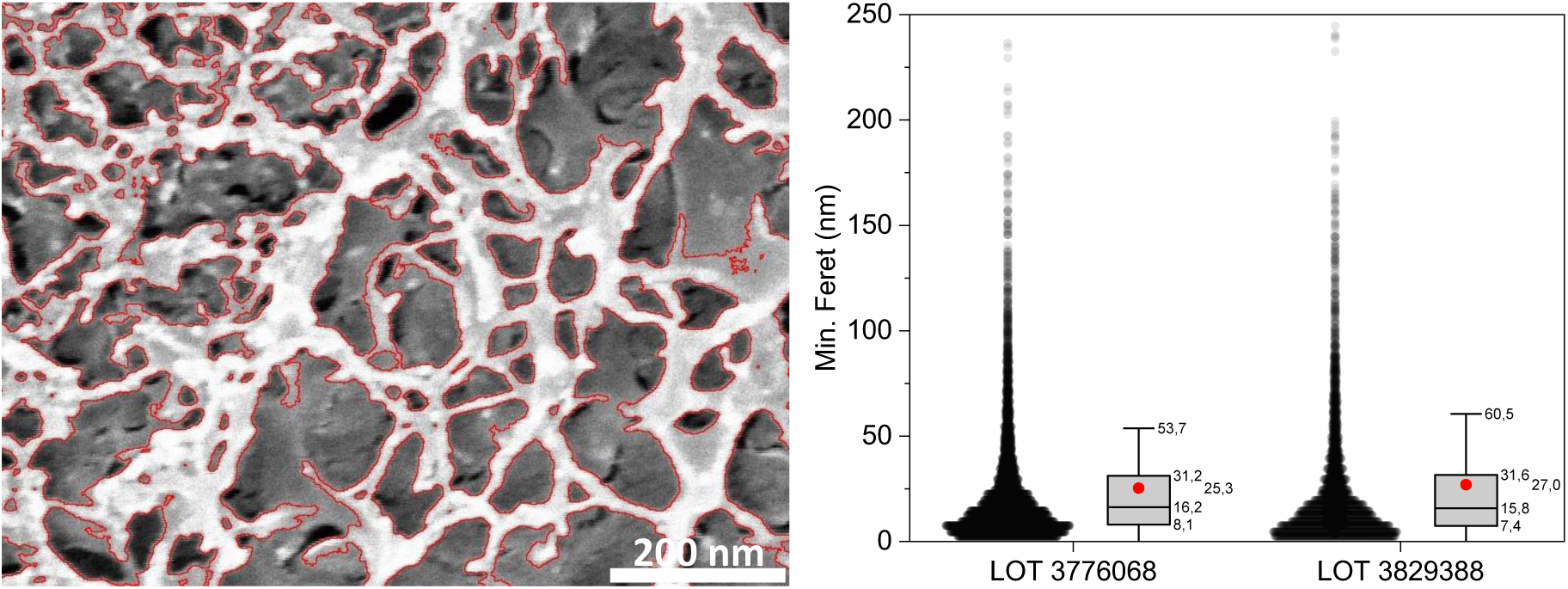
Structured analysis of SEM images. Detected pores outlined in red on original image crop of Batch-Nr. LOT 3776068 (left). Minimal Feret diameter (nm) of all pores per sample of each batch of 100 mg/mL BSM solution (right). All pore diameters are shown as individual dots and the distribution is summarized with a boxplot showing the mean as red dot, the median as black line, the 25 % and 75 % intervals as box edges and the positive single standard deviation as whisker.

### Cryo-SEM images visualize BSM network

For a further investigation of the network structure of BSM on a microscopic scale cryo-scanning electron microscopy (cryo-SEM) experiments were performed at different magnifications for the two BSM batches, in each case a homogeneous solution with a concentration of 100 mg/mL BSM dissolved in DPBS. Cryo-SEM analysis was an important tool to determine the complexity of the mucus network structure as a result of the interplay of the single components in BSM.

The images allowed the visualization of the classical web-like structure of mucus ^30, 49^. The complexity and flexibility of the mucus network and the interaction of larger and smaller mesh sizes could be identified. For both batches, areas with very small pores as well as areas with large pores were detected, i.e. a substantial structural inhomogeneity. These areas overlapped and were stacked on each other which is particularly well visible in Figure 2B; thus, forming 3D structures that can be imagined from a 2D representation such as SEM. The mucus network appeared very finely and visibly structured. There seemed to be no contamination with, for example, bacteria or cell debris. The largest visible mesh in the images was identified (20.000x magnification) and enlarged (80.000x magnification). It had a size of not more than 500 nm for both BSM batches (red boxes). More images are shown in the SI Figure S2 that confirm the general validity of the structures depicted in Figure 2.

Overall, the network structure, including the pores and pore size distribution in the mucus appear heterogeneous ^51^. This goes in line with the results from (cryo-) SEM studies from i.e. human airway mucus ^52–54^ or porcine and canine gastric mucus collections ^45, 55, 56^ Thus, BSM shows an internal network comparable to other mucus systems. The cryo-SEM preparation process was examined for its potential impact on mucus structure ^51, 53, 57^. The cryo-SEM images of the BSM displayed structures that were similar to those of mucus previously analyzed from various sources.

The pore size descriptor used was Feret’s minimum diameter, which refers to the shortest distance between two parallel tangents of a pore ^44–46^. This parameter is widely used to describe sizes and distribution of pores in SEM mucus samples. The analysis showed no significant differences between the two BSM batches in terms of their pore size. The mean values of the two batches were 25.3 and 27.0 nm and therefore only approx. 2 nm apart. The distribution of pore sizes was also similar. Accordingly, one can conclude that the mucin hydrogel structure depended only little on the choice of the BSM batch.

### Concentration Dependence of the Viscoelastic properties of BSM solutions

A central parameter that determines the viscoelasticity of hydrogels is the concentration of the network forming material. Once a critical concentration has been surpassed one typically observes a power-law increase of viscosity and shear modulus, where the exponents can vary largely as quantified for hydrophobically modified polyurethane thickeners ^58^, gelatin or guar gelled boronic acid ^59^. Accordingly, we evaluated the effect of BSM concentration by doing rheological measurements for five different concentrations: 10, 20, 60, 100, and 140 mg/mL in DPBS, all having the same pH value of 7.3, using the same batch of BSM (LOT: 38293889).

We observed an increase of the viscoelastic moduli G’ and G’’ over the investigated frequency range, this increase being clearly most marked for the more concentrated BSM solutions (60, 100, and 140 mg/mL) compared to the lower concentrated ones (10 and 20 mg/mL). For the higher concentrations significantly higher values for G’ and G’’ were observed; for instance, G’ and G’’ were around two orders of magnitude higher when comparing 100 mg/mL and 10 mg/mL BSM solutions (Figure 3A). In general, the viscous component G’’ was larger than the elastic component G’ over the investigated frequency range and the ratio G’/G’’ increased with increasing concentration (see Figure S4), which means that the elastic properties of the hydrogels get larger. In muco-obstructive lung diseases, such a large relative increase in the viscous and elastic fractions caused here by increased mucin concentration is associated with reduced transportability of mucus through the cilia and thus impaired mucus clearance ^22^. Interestingly, a dominance of G’’ over G’ can be observed in BSM, particularly at concentrations of 20 mg/mL and 100 mg/mL. In contrast, sputum from patients with CF exhibits the opposite case, with a dominance of G’ over G’’ ^26^.

**Figure 3.**
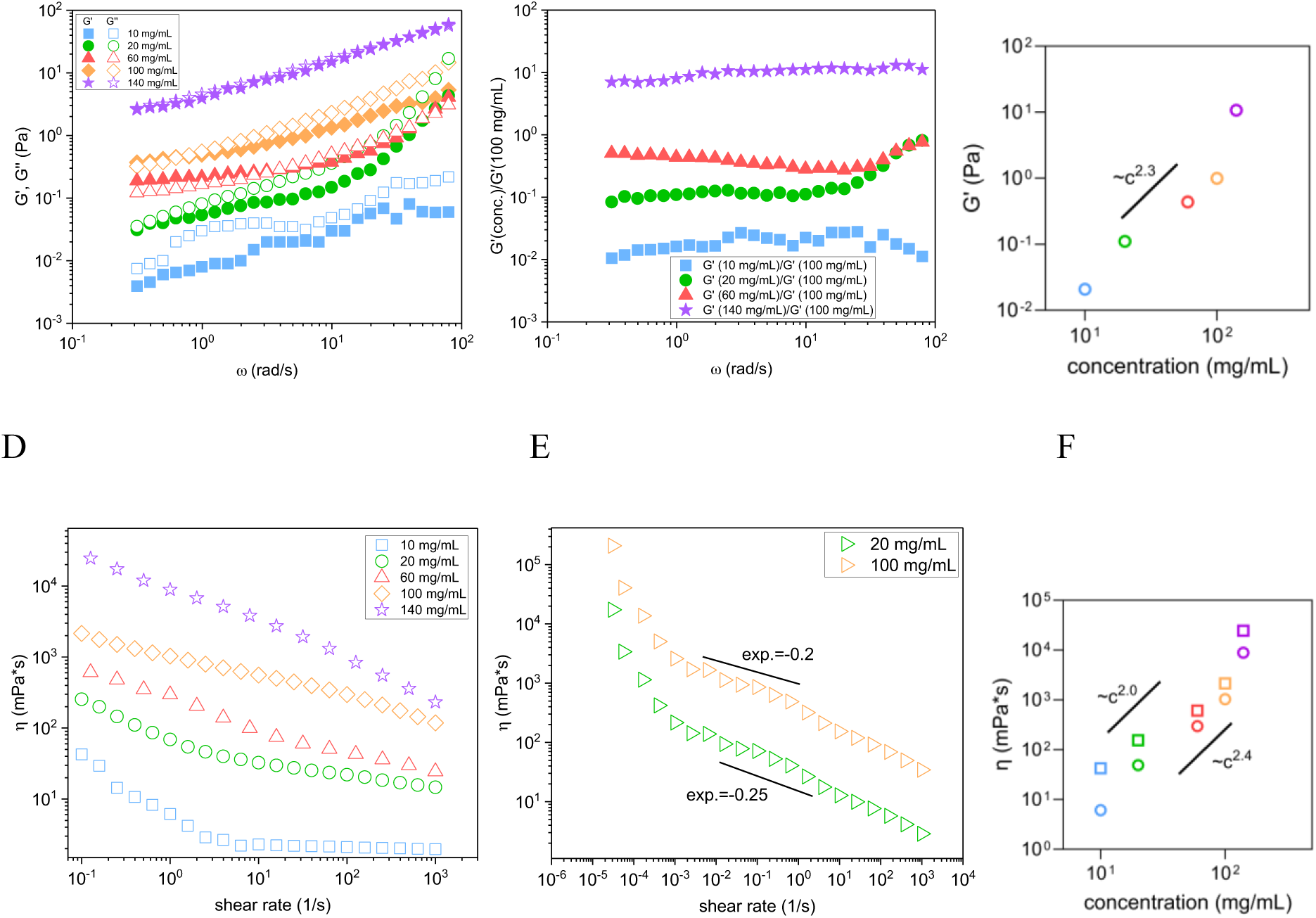
Viscoelastic properties and steady shear viscosity of different concentrated BSM solutions. Storage modulus G’ and loss modulus G’’ as function of angular frequency measured at 25 °C for BSM solutions of 10, 20, 40, 80, 100, and 140 mg/mL (deformation: 1%) (A). Ratio between storage modulus for a given concentration and that obtained for 100 mg/mL as a function of the angular frequency (B). Values of G’ as a function of concentration for frequency of 1.0 Hz (C). Data shown are mean values of n=3 measurements. Storage Steady shear viscosity as a function of shear rate; measured at 25 °C for BSM solutions of 10, 20, 40, 80, 100, and 140 mg/mL. Data shown are the mean of n=3 measurements at each concentration (D). The same measurements extended to very low shear rates (E). Viscosity at two representative shear rates (squares = 0.1 1/s and circles = 1.0 1/s) as a function of concentration (F).

When comparing the values of G’ for the different concentrations relative to G’ of 100 mg/mL concentrations one can observe an almost identical relative increase as a function of the frequency was almost identical (Figure 3B). For quantifying the concentration dependent increase of the elastic properties, we show in Figure 3C G’, taken at 1.0 Hz, as a function of concentration. One observed a power-law increase with a power law exponent of ∼2.3, a scaling law very close to that for other viscoelastic hydrogels, such as Xanthan-Al (III) gels ^60^ or chemically cross-linked polyvinyl alcohol near the gelation threshold ^61^. Theoretical work on the viscoelastic properties of entangled polymers predicts a power law with an exponent 2.0-2.3 for the plateau modulus G_0_ and an exponent of 3.4-3.7 for the zero-shear viscosity η_0_ ^62^ Apparently, BSM behaved in agreement with this model, indicating the presence of properties of entangled polymers in BSM.

Steady shear viscosity measurements showed that all BSM solutions exhibit shear-thinning behavior, which was becoming somewhat more pronounced with increasing concentration for higher shear rates (Figure 3D). Interestingly the shear rate dependence at shear rates above 10^-3^ 1/s followed a power law with a rather low exponent of 0.2-0.25 (Figure 3F), which indicates that the BSM solution was not quickly losing its viscous properties, even at rather high shear rates. This may also be important for a good functioning in biological systems.

The previously observed marked concentration dependence is similarly seen, as the 10 mg/mL BSM solution showed a significantly lower viscosity, by about a factor of ten, compared to the other concentrations for every shear rate. The differences between the viscosities of the 20 and 100 mg/mL solutions were not substantial, but they differed clearly from the viscosities of the most concentrated solution (140 mg/mL). The “zero-shear viscosity”, here approximated by the values at 0.1 or 1.0 1/s, was again decreasing with a power law with an exponent of around 2 or 2.4, respectively (Figure 3F). Interestingly, we observed that upon measuring the viscosity down to much lower shear rates below 10^-3^ 1/s a very sharp increase of viscosity was taking place. This indicated that the samples have a finite yield stress, but with a rather low value of 0.5-1 mPa, as shown in Figure S5. This is an uncommon behavior for physically entangled hydrogels, which normally show a finite maximum structural relaxation time. This matches with the general tendency seen for the BSM that the elastic properties of its hydrogels do not become much reduced upon reducing the frequency (Figure 3A).

Our finding agreed well with those of Georgiades et al. that observed for MUC5AC mucins solutions purified from porcine stomach and porcine duodenum mucin solution a power-law dependence of the shear modulus on concentration with an exponent of ∼2. The dependence of the shear viscosity on the mucin concentration in this study shows two sections, up to a concentration of 30 mg/mL the viscosity increases only slightly (exponent = 0.5) and from higher concentrations onwards with an approximately 10-fold higher exponent. The study identified a critical concentration at which the mucin solutions transition from behaving like a Newtonian liquid to forming a viscoelastic polymer network ^36^.

The consistency of elastic properties across different shear frequencies in BSM hydrogels highlights their network structure, underscoring their potential as versatile biomaterials in various applications. Interestingly, a comparable viscoelastic behavior such as the concentration dependent increase in viscoelastic moduli and shear viscosity showing similar scaling laws was observed for mucus mixtures based on polyacrylic acid (PAA) mimicking porcine intestinal mucus (PIM) ^55^.

### Detailed analysis of the rheological data of the BSM concentration series

The frequency dependent rheological data shown in Figure 3A clearly demonstrate an interesting viscoelastic behavior of the BSM hydrogels, which must result from a wide spectrum of relaxation times (i.e. cannot be described by a single relaxation time as for instance given by the Maxwell-model). In order to gain further insights into the viscoelastic properties we analyzed the different data sets with a fractional Kelvin-Voigt model (FKVM), which should be suited to describe our data as it applies to materials that are dominated by their elastic properties on longer time scales, as observed in our experimental data shown in Figure 3A. Fractional viscoelastic models are suitable for describing materials that show power-law behaviour, i.e., a slow increase of material functions over a wide frequency or time range ^63^. Detailed information on the derivation and background of the model is described in depth by Bonfanti et al. ^63^. Power law behaviour occurs in materials that exhibit a broad range of relevant microstructural length and time scales ^64^, as is the case for many biological systems ^65–68^. The FKVM, in particular, has been successfully applied to biological materials such as tissue ^69–71^, or starch gels ^72^.

In general, the FKVM consists of two spring-pots in parallel that are characterized by fractional exponents *α* and *β* and their corresponding quasi-properties *ηα* and *ηβ*. For *α* = 1, a regular viscous dashpot with viscosity *ηα* [Pa s] is retrieved and for *α* = 0, a regular elastic spring with modulus *η* [Pa] is retrieved. By definition, 1 ≥ *α* > *β* ≥ 0. For any value of *α* between 0 and 1, the quasi-property *ηα* has units of Pa s*^α^*, whose physical meaning is not straightforward, but it can generally be associated with the firmness of a material and its resistance to deformation ^75, 76^. While *ηα* and *ηβ* determine the magnitude of the response, *α* and *β* modify the model itself, since the spring-pots are becoming more spring-like or dashpot-like, respectively. For *α* = 1 and *β* = 0, the regular Kelvin-Voigt model is retrieved.

Since the BSM samples were predominantly gel-like, we fixed *β* = 0, meaning the corresponding spring-pot is a regular spring with a modulus *E* [Pa]. Fixing *β* reduces the number of fit parameters to 3, which makes the fit more robust and the interpretation easier, albeit sacrificing some fit quality. The fits with all four parameters are shown in the supporting material (Figure S10, S11) for reference. In the FKVM, the more viscous spring-pot with the higher exponent, *α*, dominates the short time-scale = high frequency behavior, while the elastic spring with *β* = 0 dominates the long time-scale = low frequency behavior. A schematic representation of the model and its constitutive equations are shown in Figure 4.

**Figure 4.**
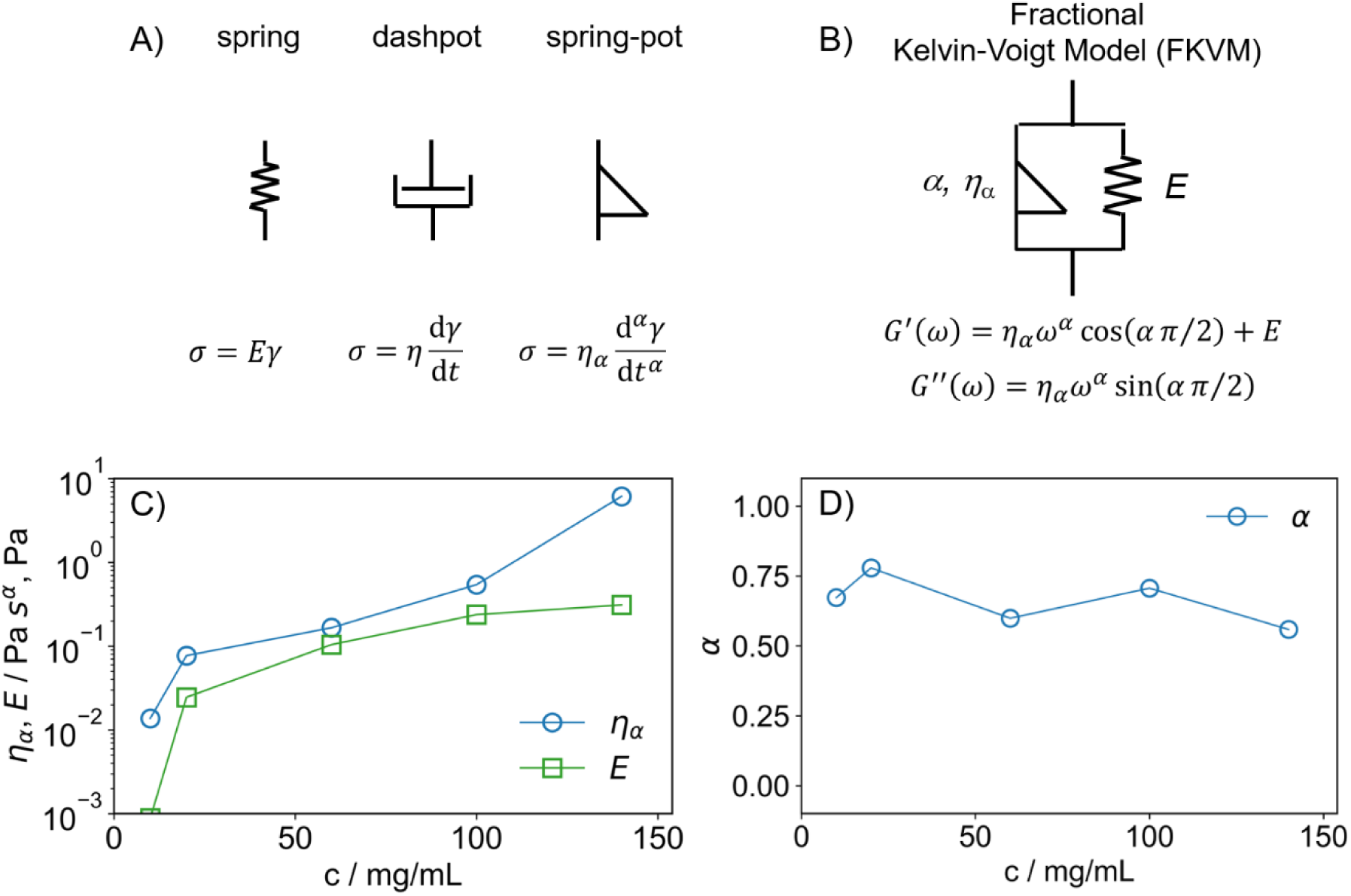
Viscoelastic models. Constitutive equations for (A) the spring, dashpot and spring-pot model elements as well as (B) the Fractional Kelvin-Voigt Model with β = 0. Parameter results for (C) the quasi-property 17_α_ (Pa*s^α^) and the modulus E and (D) the fractional exponent α from fitting the FKVM to BSM solutions of different concentrations; with γ = strain, σ = stress, t = time, G = relaxation modulus, η = viscosity (Pa*s^α,β^), *Q̄*(s) = ‘transfer function’ ^73, 74^.

The FKVM was fitted to BSM solutions at different concentrations. The results for the fit parameters are shown in Figure 4C and 4D and the fits can be found in the supporting material in Figure S10.

The fractional exponent was rather constant at a value of around *α* = 0.70, although there was a small decrease towards high concentration. This indicated that, in general, BSM solutions are well described by a model consisting of one purely elastic spring (*β* = 0) and one relatively viscous spring-pot (*α* ≈ 0.6-0.7) in parallel. It is important to point out that because *ηα* and *E* have different units, their values are not directly comparable. For both *ηα* and *E*, we first saw a steep increase from 10 to 20 mg/mL, followed by a very flat increase until 100 mg/mL. In this rather wide concentration regime, surprisingly, the viscoelastic properties hardly changed, which was very different to what is normally observed in viscoelastic polymer solutions ^77^. This could be an important aspect for its *in vivo* performance, as apparently its general viscoelastic properties change only rather little over an extended concentration regime. Going to 140 mg/mL, *ηα* increased much more strongly than *E*, indicating that the relative importance of the viscous properties versus the elastic properties became larger.

### Effect of additives

After the establishment of concentration dependence of the viscoelastic properties of BSM we investigated how they become modified by the presence of different additives. To do so, we added different salts, proteins, or DNA to the mucus samples; all components that may be present to a larger or lesser extent in biological mucin samples and that are relevant for its function.

#### Sodium chloride

In our initial investigation, we examined the impact of NaCl, as sodium ions could potentially alter the hydration state of mucus and thus may cause a change in its viscoelastic properties ^78, 79^. For that purpose, we performed rheological measurements with 100 mg/mL BSM solutions with four different NaCl concentrations, where one may expect the main effect arising from an increase of the ionic strength of the solution and only a lesser effect from specific binding of the ions to the mucins. The resulting BSM solutions had sodium chloride concentrations of 0.2, 0.4, 2.2 and 5.2 M. A 100 mg/mL BSM solution dissolved in DPBS contains already 0.15 M sodium chloride, resulting from manufacturing of the commercial BSM. The pH value of all solutions was around 6.9.

Previous experiments on mucins purified from tracheobronchial mucus secretions from CF patients and healthy subjects were suggested to be present as a less extended flexible coil structure in the presence of NaCl ^80^. This study aimed to investigate the impact of NaCl concentration on mucin aggregation and the resulting viscoelastic properties of CF lung mucus secretions, which are essential for effective mucus clearance due to its transport capacity. Structural changes in the internal architecture of the mucins due to the addition of sodium may also occur in the NaCl containing BSM solutions which then should also affect the rheological behavior. It seems that a small increase in salinity can enhance the mucus clearance from the respiratory tract (tested here up to 90mM) ^81^, which should be related to the reduction of the viscoelastic properties.

According to the rheological measurements shown in Figure 5 A-C, the addition of NaCl to a 100 mg/mL BSM solution led to a generically similar increase in the viscoelastic moduli G’ and G’’ with increasing frequency for all five NaCl concentrations. The absolute values first decreased with the concentration of added NaCl, but then increased again a very high NaCl concentrations (Figure 5A). Here the solution with 0.2 M NaCl in a 100 mg/mL BSM solution had lower values of G’ and G’’ value at low frequencies by almost a factor five compared to the solution with 0.15 M NaCl, which was an interesting effect for the addition of just 50 mM NaCl.

The phase angle (Figure 5B) showed that the relative relevance of the elastic component becomes first reduced with increasing NaCl concentration and then became more significant again for the two highest NaCl concentrations (see also Figure S6). However, here it has to be noted that in these samples with 2.2 and 5.2 M NaCl almost all the water molecules are directly bound to the ions and therefore one may effectively see a more concentrated BSM network. In addition, for most viscoelastic polymer solutions it is observed that with increasing frequency the relative contribution of the elastic component becomes more important ^82^, but for the different BSM solutions an opposite behavior was noted. This is an intriguing behavior that showed that for higher frequencies the ability of the BSM hydrogel to dissipate energy increased compared to its ability to store energy elastically. These findings suggested that the primary effect arises from an increase in ionic strength rather than specific ion binding to mucins.

For all samples shear thinning behavior was observed in the steady shear viscosity experiments (Figure 5C). The 0.2 M, 0.4 M and 2.2 M sodium BSM solutions showed similar curves, starting from just under a 1000 mPa·s for low shear rates and then decreased to 100 mPa·s at higher shear rates. In contrast, the highest NaCl concentration of 5.2 M showed a viscosity value of higher than 1000 mPa·s, decreased more rapidly with increasing shear rate, and reached the same values as the other sodium BSM solutions at a shear rate of around 2 1/s.

**Figure 5.**
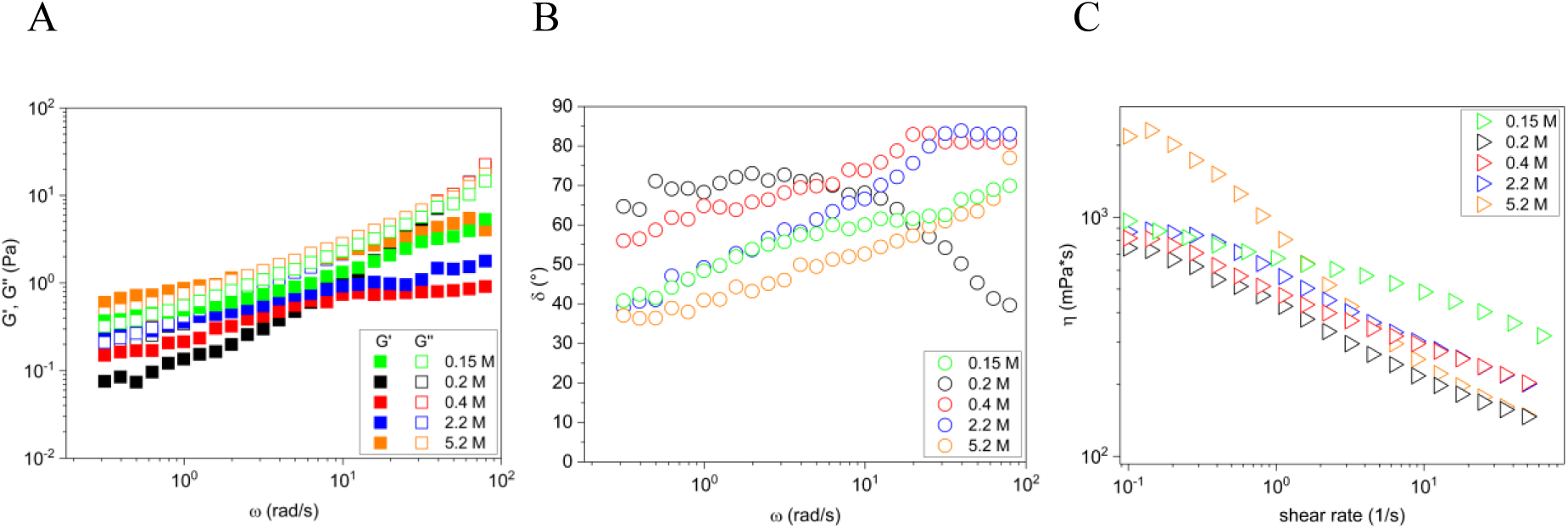
Viscoelastic properties of BSM solutions with added NaCl. Storage modulus G’ and loss modulus G’’ and corresponding phase angle δ of 100 mg/mL BSM solutions with 0.15 M (baseline), 0.2 M, 0.4 M, 2.2 M and 5.2 M NaCl were measured as function of angular frequency (rad/s) at 25 °C (A and B). With a new aliquot of each solution the steady shear viscosity as a function of shear rate was measured (C). Data are shown as mean of n=3 measurements of each concentration.

#### Calcium chloride

To investigate the effect of calcium ions (as CaCl_2_), which is known to have an important influence on mucus network properties ^1, 9, 21^, we studied 100 mg/mL BSM solutions with different CaCl_2_ concentrations. The 100 mg/mL BSM solution contained already an initial concentration of 0.1458 mg/mL calcium (= 1.0 mM) (see SI, ICP-OES) and it was dissolved in a HEPES buffer that contained no additional calcium. The final calcium concentrations of the solutions were 1.0, 2.7 mM, 6.0 mM, 11.0 mM, 21.0 mM, 51.0 mM and 75.0 mM. The pH value of all solutions was around 6.7.

The 100 mg/mL BSM solution was experimentally determined to have a total protein content of approximately 85 mg/mL (refer to Methods section), with around 10% of this amount constituting sialic acid in free and glycosidicically bound form within the mucins. This estimation is based on experimental data from previous studies ^83, 84^, corresponding to about 8.5 mg/mL or 25-30 mM of sialic acid. At this sialic acid concentration, crosslinking effects are anticipated with the addition of 10-20 mM calcium, as it was observed in the experiments. The calcium added to the BSM solution is likely to bind to the negatively charged acid groups of the sialic acid, along with other acidic groups in mucin backbone or calcium-binding sites, potentially enhancing intermolecular interactions. With higher calcium concentrations, it is expected that all carboxylic groups would become complexed, leading to reduced crosslinking. This aligns with the observation that at the highest calcium concentration of 75 mmol/L (equivalent to 11 mg/mL), there was a noticeable decrease in the viscoelastic properties.

The addition of CaCl_2_ led to substantial changes of the rheological properties already at rather low CaCl_2_ concentrations (Figure 5 D-F), different to the situation observed for NaCl addition (Figure 5 A-C). A continuous increase of the viscoelastic moduli was seen with increasing CaCl_2_ concentration, where the maximum was reached for the 51 mM sample. Interestingly for the sample with the highest CaCl_2_ concentration of 75 mM, a concentration far above that found in nature, the moduli became reduced again by about a factor 10 and were then lower than for the case without added CaCl_2_ (Figure 5D). This reduction of the viscoelastic properties is likely related to the compaction of mucins (e.g. MUC5B) that has been reported in the presence of excess Ca^2+^ ions ^21, 85^. We detected several crossover points for these low concentrations, which is better seen in the phase angle depicted in Figure 5E. When performing frequency sweeps up and down, we noted identical curves, indicating reversible deformation of the samples.

For the phase angle (Figure 5E) rather constant values were observed with a slight increase with increasing frequency. The sample with 51 mM CaCl_2_ not only had the highest moduli but also the elastic part was most pronounced. The samples with low CaCl_2_ concentration (1.0 and 2.7 mM) showed G’ and G’’ values around 0.5 Pa increasing up to 5 Pa with increasing frequency, i.e., somewhat more marked viscous properties (phase angle increases to 65° at higher frequencies).

The steady shear viscosity experiments with the same CaCl_2_ BSM solutions showed a similar shear thinning behavior like for the pure BSM solutions, but at low shear rates the viscosity was significantly higher compared with pure BSM solution. We observed the highest viscosities for the 21.0 mM and 51.0 mM calcium BSM solutions at low shear rates, where they were higher by a factor 100. However, they drastically decreased with increasing shear rate; reaching at a shear rate of around 1000 1/s the same values of about 100 mPa·s, irrespective of CaCl_2_ concentration. This means the shear-thinning effect was much more pronounced for the samples with 21.0 and 51.0 mM CaCl_2_ and these samples were also more resistant with their viscous properties against shear forces. The samples with the highest added CaCl_2_ concentration showed a much less pronounced decrease in viscosity with increasing shear rate, with a lower slope. The viscosity of the 75.0 mM calcium-BSM-solution appeared in the same range then the viscosity of the low (2.7 mM) or no (1.0 mM) added calcium.

In summary, it can be stated that the addition of concentrations of about 10 to 50 mM CaCl_2_ enhances the gel properties of the BSM system, where this effect was rather sensitive in its extent to the CaCl_2_ concentration.

The sensitivity of mucus systems to calcium has been observed in various solutions, such as PGM and MUC5B solutions purified from human saliva. An increase in viscoelastic parameters was found with the addition of calcium concentrations of approximately 10 mM. This increase was ascribed to stronger entanglement of the mucins and the resulting more compact network structure ^9, 21^. For example, in CF the deficiency of CFTR-mediated bicarbonate transport results in more free calcium, which can contribute to mucin complexation ^85^.

**Figure 5.**
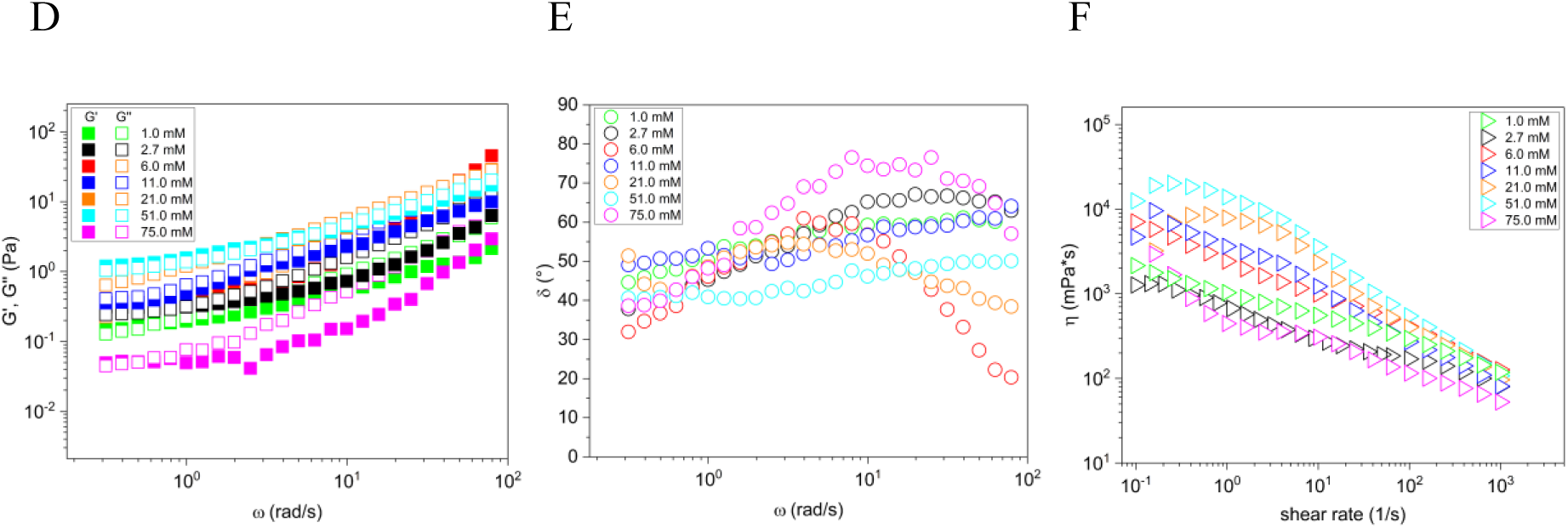
Viscoelastic properties of BSM solutions with added CaCl_2_. Storage modulus G’ and loss modulus G’’ and corresponding phase angle δ of 100 mg/mL BSM solutions with 1.0, 2.7, 6.0, 11.0, 21.0, 51.0, and 75.0 mM CaCl_2_ as function of angular frequency (rad/s) at 25.0 °C (D and E). With a new aliquot of each solution the steady shear viscosity as a function of shear rate was measured (F). Data are shown as mean of n=3 measurements of each concentration.

#### Lysozyme

Apart from small ions, native mucus is surrounded by a variety of non-mucin proteins, hence their contribution to the viscoelastic characteristics of mucus needs to be understood. For this purpose, we studied the effect of lysoyzme acting here as a model protein on the viscoelastic response of BSM.

Lysozyme, a key component in various biological fluids, plays a crucial role in mucosal defense due to its antimicrobial properties, achieved through the hydrolysis of β–(1,4)–glycosidic bonds in the peptidoglycan of pathogenic microorganisms’ cell walls ^86^, and its anti-inflammatory effects^87^. Its interaction with mucins, such as the ability to form complexes with porcine gastric mucin (PGM) through various binding types ^86^, underscores its significance in mucosal biology and prompted our investigation into its effects on BSM. It is worth noting that the BSM did not inhibit lysozyme activity, which contrasts with the PGM that did so under certain conditions ^88^.

Here, 100 mg/mL BSM solutions with 1.0, 2.0 or 10.0 mg/mL lysozyme were measured at a pH-value of 7.2. At this pH lysozyme with an isoelectric point of 11, is fully positively charged (∼8 positive charges). Following, lysozyme, acting as a cross-linking agent, strong electrostatic intereactions between negatively charged mucins and positvely charged lysozyme are taking place ^86^.

The data given in Figure 5 G-I showed for the lower lysozyme concentrations, that the rheological moduli did not change much compared to the situation without added lysozyme, and the loss modulus G’’ was larger than the storage modulus G’ within the investigated frequency range, resulting in a constant phase angle at 70° for both lysozyme concentrations. For 10.0 mg/mL lysozyme the moduli were much larger (Figure 5G) and here G’ was about the same as G’’, resulting in a much lower phase angle δ (Figure 5H). Interestingly, this sample differed even more with respect to the viscosity, as it had a much higher viscosity, which was also visible by eye. The viscosity decreased to the level observed in low concentration solutions at high shear rates (100 mPas) with a rather strong power law with an exponent of -0.927. The much more marked viscosity of the BSM sample with 10.0 mg/ml lysozyme added was not due to the relatively high concentration of lysozyme, as shown by a comparative measurement of a sample with the same lysozyme concentration but without BSM, which shows a 100-1000 times lower viscosity (Figure 5I). The cause is likely attributed to electrostatic interactions between the positively charged groups on the lysozyme and the negatively charged sites on the mucin molecules ^86, 89^. This observation was also noted in sputum from patients with chronic obstructive pulmonary disease (COPD) to which lysozyme was added ^89^.

**Figure 5.**
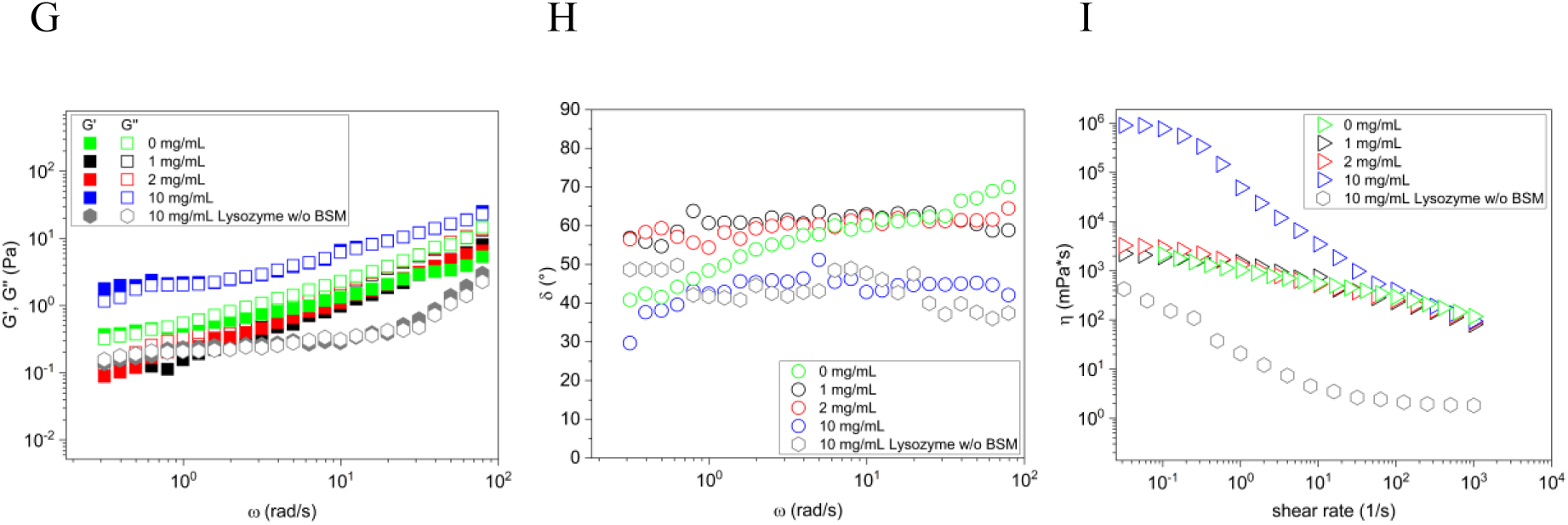
Viscoelastic properties of BSM solutions with added Lysozyme. Storage modulus G’ and loss modulus G’’ and corresponding phase angle δ of 100 mg/mL BSM solutions with 0, 1.0, 2.0, and 10.0 mg/mL Lysozyme as well as a 10 mg/mL Lysozyme solution without BSM as function of angular frequency (rad/s) at 25 °C (G and H). With a new aliquot of each solution the steady shear viscosity as a function of shear rate was measured (I). Data are shown as mean of n=3 measurements of each concentration.

#### DNA

We investigated the effect of DNA on mucus viscoelastic properties, which are known to be influenced by DNA ^90^ e.g. in the context of neutrophilic granulocyte activation, which burst and release DNA during inflammation. Especially, in chronic muco-obstructive lung diseases such as cystic fibrosis ^91, 92^ therapeutic strategies are based on DNAse supported mucin dilution ^90^. We studied 100 mg/mL BSM solutions with varying concentrations of DNA. The determination of the initial DNA content in a 100 mg/mL BSM solution showed 1.2124 mg/mL DNA with a standard deviation of ± 0.000743 mg/mL. In small steps of 5 mg/mL we increased the DNA concentration up to 20 mg/mL. The pH value of the solutions was around 7.07.

The solution with the smallest amount of DNA (baseline 1.21 mg/mL) had the highest values for the viscoelastic moduli with a prominent difference of G’’ higher than G’. The DNA concentration of 15 mg/mL came close to these values, but with an increasing DNA concentration the viscoelastic moduli decreased, and there were nearly no differences between the G’ and G’’ (Figure 5J). All the solutions showed shear thinning behavior with nearly the same power law exponent of around -0.3 (Figure 5L). Again, the 15 mg/mL concentrated DNA solution had the highest values for viscosity, but approached very quickly, already at a 1 1/s shear rate to the other solutions with added DNA.

**Figure 5.**
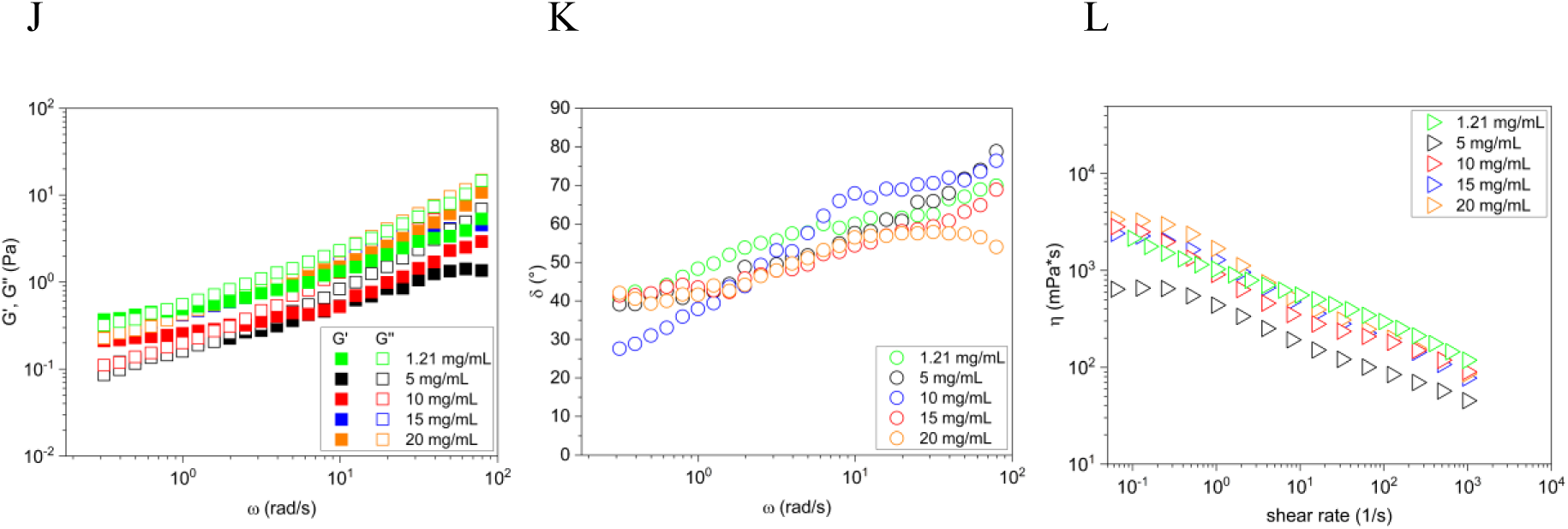
Viscoelastic properties of BSM solutions with added DNA. Storage modulus G’ and loss modulus G’’ and corresponding phase angle δ of 100 mg/mL BSM solutions with 1.21 mg/mL (baseline), 5.0 mg/mL, 10.0 mg/mL, 15.0 mg/mL and 20 mg/mL DNA were measured as function of angular frequency (rad/s) at 25 °C (J and K). With a new aliquot of each solution the steady shear viscosity as a function of shear rate was measured (L). Data are shown as mean of n=3 measurements of each concentration.

Initially, it was expected that the viscoelastic moduli would increase with increasing DNA concentration in the BSM. Interestingly, as the DNA concentration increased, the viscoelastic moduli decreased, and the difference between G’ and G’’ narrowed, suggesting that higher DNA concentrations alter the viscoelastic balance in BSM solutions. It may be hypothesized that the DNA concentrations added were already too high to interact rheologically visible with the mucus structure. This trend, along with the observation of consistent shear-thinning behavior across different DNA concentrations, highlights the complex interaction between DNA and mucin molecules, underscoring the importance of DNA concentration in modulating the rheological properties of mucus-like systems.

Lehr et al. pointed out the importance of DNA in mucus models ^10^. They compared the rheological behavior of various mucus surrogates with human native pulmonary mucus, taking into account the elastic-dominant behavior when selecting components for the mucus model. This partially simulates the rheological behavior of human native pulmonary mucus. To approximate the viscoelastic properties of human airway mucus with the BSM, our study suggests making specific adjustments, such as adapting the elastic and viscous fractions within certain limits using the described calcium and DNA concentrations, which would enable predictions to be made using the FKVM.

#### Detailed analysis of the rheological data of the BSM solutions containing additives

We once more utilized the Fractional Kelvin-Voigt Model to analyze the rheological behavior of the previously described BSM solutions with additives. The results for the fit parameters are shown in Figure 6, the corresponding fits can be found in the supporting material in Figure S10.

**Figure 6.**
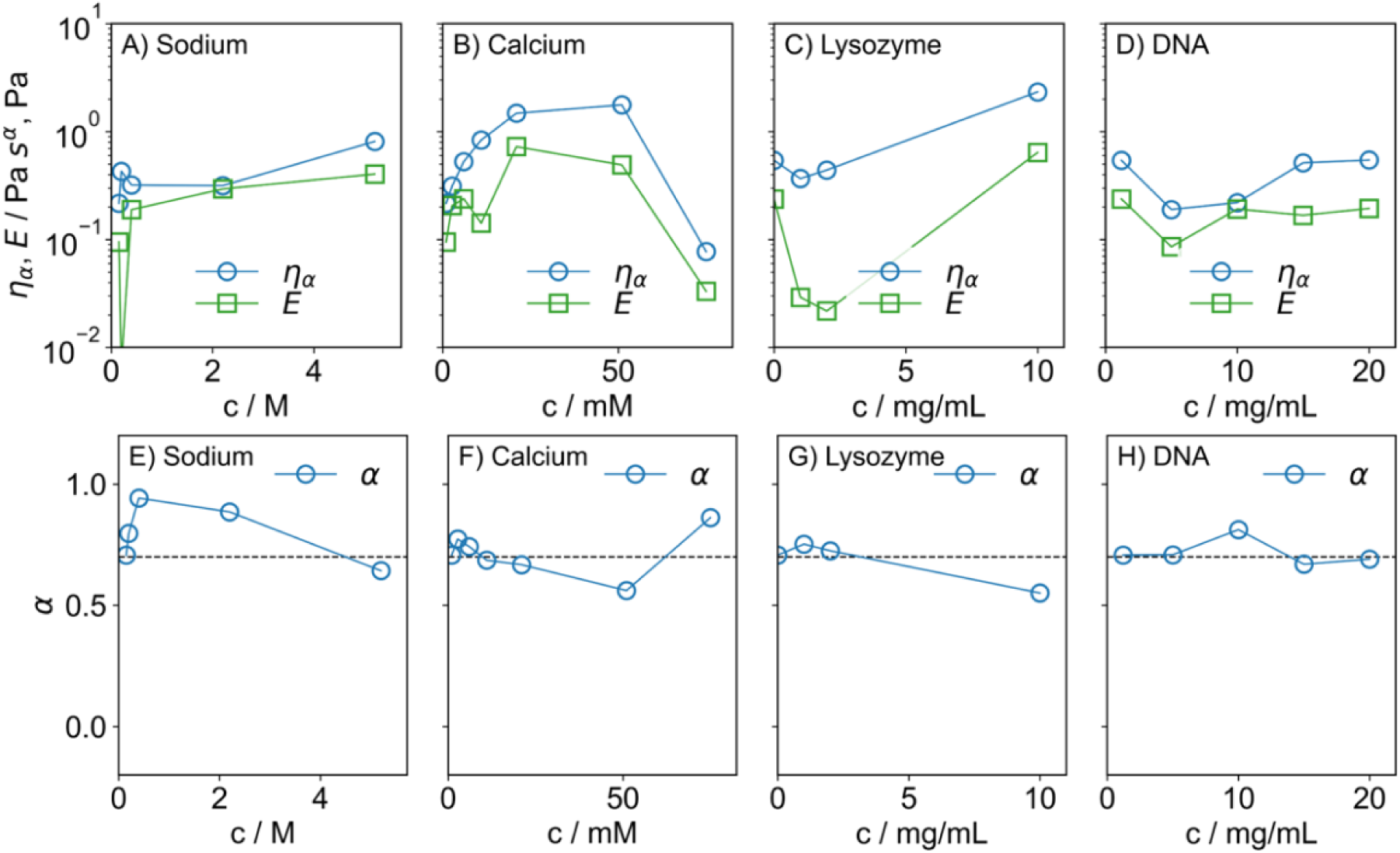
Fitting parameters obtained from FKVM. Variation of FKVM parameters as a function of concentration of additives. A)-D) Variation of the quasi-property 17_α_ and the modulus E, E)-H) Variation of the fractional exponent α. The dashed line indicates a value of α = 0.7.

In general, variation of the model parameters with the concentration of additives is rather small. Upon further examination, several interesting features merit attention. Similarly to what was seen for the BSM concentration dependence discussed above, the fractional exponent *α* remained close to a value of 0.7, which is indicated by dashed black lines in Figure 6 E-H. Systematic devations were only seen for sodium, where *α* first sharply increased between 0 and 0.4 M to a value of 0.94, before decreasing again to 0.64.

For sodium, there was a slight increase of both *η_α_* and *E* towards higher concentrations. Although the influence is rather small, NaCl thus tended to strengthen the network structure, most likely through electrostatic interaction with negatively charged parts of mucins or other proteins that can be found in BSM. This could result in changes to the structure of the mucin, as well as an effectively higher concentration of mucin, due to the increased hydration of the ions^80^.

The BSM solutions showed a much more pronounced dependence on added Ca^2+^ compared to the addition of Na^+^. With increasing calcium concentration, both *η_α_* and *E* first increased before reaching a maximum between 21 and 51 mM. For the highest calcium concentration, 75 mM, we saw a marked decrease of *17α* and *E*, which again mirrors the known compaction of MUC5B, already described above.

A concentration dependence was noted for the addition of lysozyme, where *E* first decreased up to 2 mg/mL before increasing again for the highest concentration at 10 mg/mL. The elastic properties were first weakened and then strengthened by the presence of lysozyme. *17α* showed only a slight decrease before significantly increasing at 10 mg/mL.

The addition of DNA to a BSM solution did not seem to have any effect on the viscoelastic properties of BSM. The FKVM fitted data align well with the experimental data.

Overall, the rheological properties of BSM solution seem to be largely unaffected by the presence of sodium, lyoszyme and DNA. Calcium was an exception, which has a very marked influence on the FKVM parameters at much lower concentration in the mM regime.

The FKVM allowed us to completely describe the complex viscoelastic behavior of mucus, while simpler models could not. This is mainly due to the wide spectrum of relaxation times inherent in BSM. The FKVM, particularly suited for materials exhibiting power-law behavior and a dominance of elastic properties over longer time scales, was an ideal choice for our analysis, as it effectively describes materials with a broad range of microstructural length and time scales, typical of many biological systems including BSM. This model allowed us to capture the nuanced viscoelastic behavior of BSM using a minimal set of model parameters, distinguishing between the interplay of its viscous and elastic components over different frequency ranges for the behavior at increasing BSM concentrations and with additives.

### Proteome

Commercial mucins underwent several purification steps before it is sold as lyophilized powder. We surmised several other proteins, and DNA inside of commercial BSM, and therefore conducted proteomics and DNA content analysis. Overall, we identified and quantified 2645 proteins in the replicates (Figure 7A). Three of these 2645 proteins were mucins: MUC19, and two different submaxillary mucin fragments that we numbered with sequence 1 (S1, Uniprot ID Q9N1P0) and sequence 2 (S2, Uniprot ID O62672) as they have the same gene symbol BSM1. All three proteins have partially overlapping sequences but unique peptides were identified for each of them (Figure 7B). These mucins determined in this examination highlighted the diverstiy within the protein composition of BSM.

**Figure 7.**
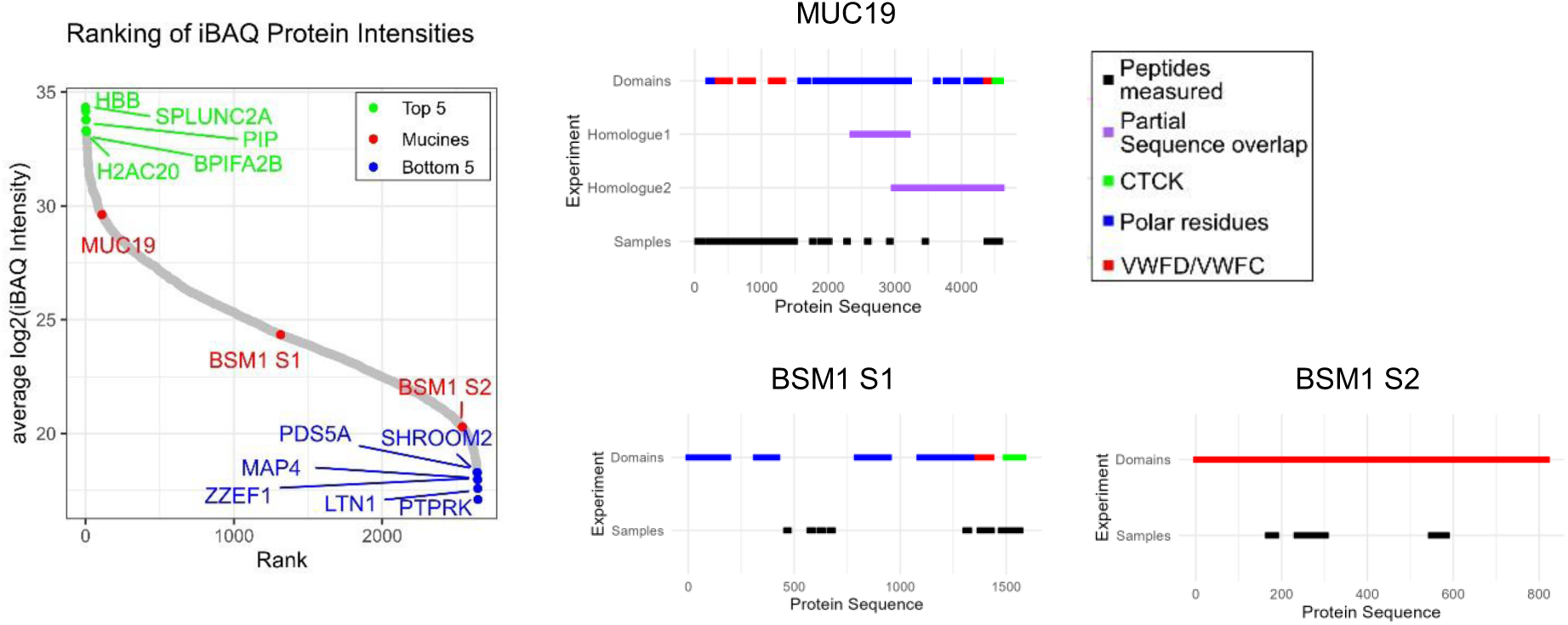
Ranking of quantified proteins according to their average log2 iBAQ normalised intensity. The top 5 and bottom 5 proteins are highlighted in addition to the quantified mucines MUC19, BSM1 S1 (Uniprot ID Q9N1P0) and BSM1 S2 (Uniprot ID O62672) (A). Peptides measured in the three technical replicates for each mucin are highlighted together with overlapping sequences (purple), and the three different domains: carboxyl-terminal cystine knot (CTCK) (green), von Willebrand Factor type D and C (VWFD/VWFC) domains and the polar residue rich domain (proline, theronine, and sering (PTS)) as annotated on Uniprot.

## CONCLUSIONS

In conclusion, the comprehensive investigation of Bovine Submaxillary Mucin (BSM) in this study provides valuable insights into its biomechanical behavior in the context of several physiologically relevant stimuli and presents a multiparametric function to describe viscoelastic responses of this biomaterial. Our findings reveal that the viscoelastic properties of BSM only somewhat increase with material concentration, which is important for robustness in biological systems. Addition of calcium ions markedly enhances viscoelasticity of BSM, while changes in ionic strength resulting from sodium chloride concentration have minimal effects. The application of a mucus analysis strategy using the Fractional Kelvin-Voigt Model captures the complete rheological behavior of BSM, utilizing a minimal set of model parameters. Cryo-scanning electron microscopy effectively revealed the intricate internal network structure of BSM, facilitating a detailed analysis of pore sizes, while complementary proteome analysis provided a deeper understanding of the BSM composition, which identified MUC19 as the predominant gel-forming mucin in BSM.

Our findings and modifications to BSM offer a solid foundation for developing targeted human airway mucus models, which can be adjusted to meet specific research requirements. Ribbeck et al. have previously discussed the challenges associated with this, in their research on mucus surrogates derived from tissue-specific mucins to simulate various types of human mucus^16^. Although it is feasible to directly tailor the physico-chemical properties of these models to match human mucus sources is feasible under certain conditions, it remains a complex task, particularly when considering the full spectrum of mucus characteristics across various length scales. However, the modifications to BSM explored in our study provide a robust foundation for developing targeted human airway mucus models that can be fine-tuned to meet specific research requirements. This approach allows for more accurate simulations of human mucus behavior, enhancing our understanding and potential treatment strategies for various respiratory conditions.

In comparison to other studies on mucins and mucus, the BSM rheological behavior BSM makes it an important mucus source to study responses to different physico-chemical situations relevant for mucus dynamics under physiological and pathophysiological conditions. Our analytic FKVM term provides researchers from various fields with a fundamental basis to study this biomaterial as a surrogate for human lung mucus in greater detail.

## ASSOCIATED CONTENT

### Supporting Information

Additional experimental details, materials, and methods, as well as complementary data (PDF).

## AUTHOR INFORMATION

### Author Contributions

Conception and design of the study: HR, MAM, DL, MG

Acquisition, analysis, and interpretation of the data: HR, RFS, LFW, KF, YK, PM, MAM, DL, MG

Drafting the article or revising it critically for important intellectual content: HR, RFS, LFW, KF, YK, PM, MAM, DL, MG

All authors contributed to the article and approved the submitted version.

## ACKNOWLEDGMENT

We acknowledge the support by the German Research Foundation. We would like to express our gratitude to Dr. Brigitte Tiersch for taking the cryo-SEM images, and to Professor Joachim Koetz from the Colloid Chemistry Department, Institute of Chemistry, University of Potsdam. Special thanks to Jana Lutzki, Stranski-Laboratorium für Physikalische und Theoretische Chemie, Technische Universität Berlin, and Dr. Wiebke Fischer, Institute for Chemistry and Biochemistry, Freie Universität Berlin.

