## Supplementary Information for "Comprehensive characterization of the viscoelastic properties of Bovine Submaxillary Mucin (BSM) hydrogels and the effect of additives"

#### Methods

##### Measurement of the calcium and sodium content in BSM with Inductively Coupled Plasma - Optical Emission Spectrometry (ICP-OES)

BSM solutions with concentrations of 1 mg/mL (five repetitions) and 5 mg/mL (five repetitions) were prepared using a solvent mixture of distilled water, nitric acid 65 % (Merck KgaA, Suprapur, CAS-number: 7697-37-2) and hydrogen peroxide 30 % (ChemSolute, Th. Geyer, Nr. 476.1011) in a ratio of 5:3:1. The solutions were stirred as stated above. The solutions were digested by a Microwave digestions system (Discover SP-D, CEM) with a fixed heating and cooling protocol; first heating up to 160 °C at 17 bars for 20 min and then cooling down to 70 °C at 17 bar for 10 min. Afterwards the solutions and reference solutions of three different concentrations 1.0, 10.0, and 100.0 mg/L (Multielement Standard Solution 5 for ICP, Merck KGaA, 54704) were analyzed for their calcium and sodium content by using Optical Emission Spectrometry with Inductively Coupled Plasma (ICP-OES) with a Varian ICP-OES 715 ES spectrometer. The parameters of the measurement were the following: plasma gas 16.5 L/min (argon), assistance gas 1.5 L/min (argon), atomizing pressure 200 kPa, observation height 10 mm, stabilization time 25 s, time for sample intake 35 s, measurement time 10 s, pumping speed 15 upm, flushing time 60 s. The determination of the wavelength used for the evaluation of the data for calcium and sodium was made based on the information in DIN EN 16943 - 2017-07 and DIN EN ISO 11885 - 2009-09; wavelength calcium 317.933 nm and sodium 588.995 nm.

##### Baseline sodium and calcium concentrations of BSM determined via ICP-OES

The sodium concentration in BSM solutions (100 mg/mL) of the two batches dissolved in water was determined by five independent measurements at a sodium-specific wavelength of 588.995. The calculated sodium concentration is 1.1263 ± 0.0005 mg/mL.

In the same way, the calcium concentration was determined from five independent measurements at a wavelength of 317.933. It is 0.1458 ± 0.0002 mg/mL for 100 mg/mL BSM solutions of the two batches (dissolved in water).

##### Addition of salts, DNA and lysozyme to a 100 mg/mL BSM solution

The production of the four BSM sodium solutions proceeded as follows: DPBS buffer, containing 8 mg/mL sodium, was added to each 100 mg BSM (containing 1.12 mg/mL sodium, which was determined from ICP-OES measurements) and each 3 mg, 11 mg, 117 mg and 292 mg sodium chloride (Sigma-Aldrich, ≥99.5 %, S9888). This resulted in four BSM solutions with the following sodium chloride concentrations: 12 mg/mL (= 0.2 mol/L), 21 mg/mL (= 0.4 mol/L), 126 mg/mL (= 2.2 mol/L) and 301 mg/mL (= 5.2 mol/L). A 100 mg/mL BSM solution dissolved in DPBS was taken as baseline reference having a 9 mg/mL sodium chloride concentration (= 0.15 mol/L).

The production of the six BSM calcium chloride solutions proceeded as follows: HEPES buffer was added to 100 mg BSM (containing 0.15 mg/mL calcium, which was determined from ICP-OES measurements) and each 0.22 mg, 0.74 mg, 1.5 mg, 3.0 mg, 7.0 mg and 11.0 mg calcium chloride (via calcium chloride dihydrate, Merck KGaA, ≥99.0 %, C7902). This resulted in six BSM solutions with the following final calcium chloride concentrations: 0.4 mg/mL (= 2.7 mmol/L), 0.9 mg/mL (= 6 mmol/L), 1.6 mg/mL (= 11 mmol/L), 3.1 mg/mL (= 21 mmol/L), 7.5 mg/mL (= 51 mmol/L) and 11 mg/mL (= 75 mmol/L). A 100 mg/mL BSM solution dissolved in HEPES was taken as baseline reference containing 0.15 mg/mL calcium (= 1 mmol/L).

##### Quantitative analysis of Cryo-Scanning Electron Microscopy (cryo-SEM) images

The pore size distributions of the mucus samples were determined by automatic analysis of cryo-SEM images with a self-written Fiji [1] macro. The macro can be used to analyze single images and image stacks. First an image or stack must be opened in Fiji and then a region of interest (ROI) must be selected (e.g., whole image excluding scale bar). The macro can then be executed by pressing *Run*.

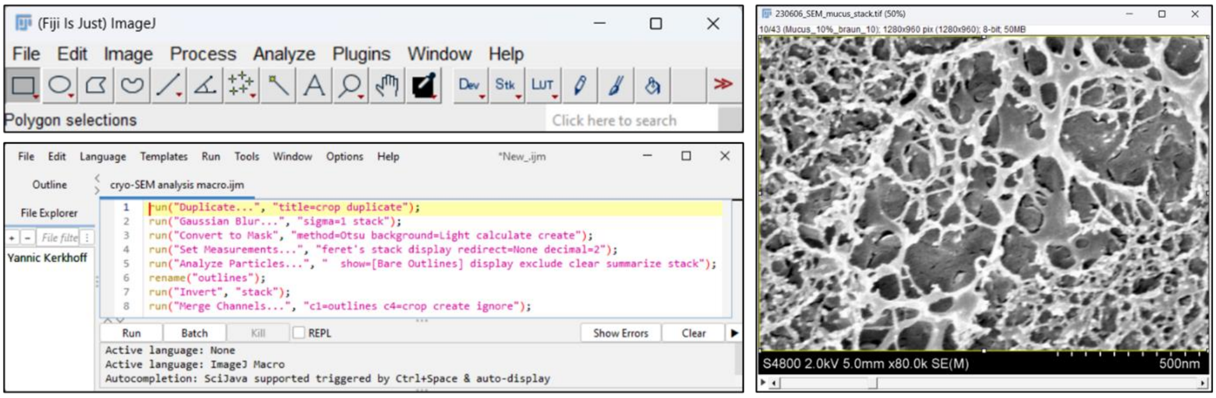

***Figure S1A:*** *The Fiji action bar with the opened analysis macro (left) and the image stack containing cryo-SEM images with a selected ROI (yellow box) excluding the scale bar at the bottom.*

The ROI will be duplicated and renamed “crop” and a Gaussian filter with a pixel radius of 1 will be applied to reduce noise [2]. The images are then binarized by an Otsu-based threshold [3] to separate the pixels into two classes (pores, fibers) with minimized inter-class variance. The binary images named “MASK_crop” are then analyzed (Area, Feret) with the Particle Analyzer included in Fiji. The analyzed pores are outlined where pores touching the edges of the images are not included into the analysis to avoid a bias origin from incomplete pores.

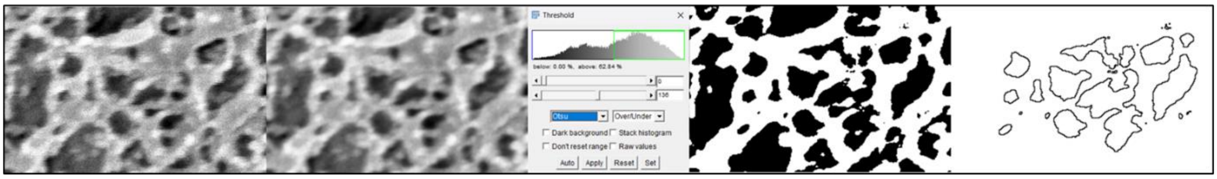


***Figure S1B:*** *Representation of the basic image processing and analysis steps. From left to right: Crop of one original cryo-SEM image of Batch-Nr. LOT 3776068. A Gaussian filter with a pixel radius of 1 is applied to reduce noise. An Otsu-based threshold transforms the image into a binary image, with black pores and white fibers. The pores not touching the edges are detected and outlined by the Particle Analyzer.*

The outlines of the detected pores are overlayed in red onto the original images (“Composite”) for manual inspection of the detection quality. When the macro is finished, the original images are retained, as well as the binary images, which can be saved for further analysis (e.g., with exclusion of small objects). Two data tables are created. The *Results* table contains the values of all single pores. The *Summary* table contains averaged values of all pores per image for quick comparison of different images. The values in the data tables are in pixel units and must be multiplicated with the respective pixel size (e.g., in nm) to get the correct pore size distributions. As a key marker for the pore diameter the minimum Feret is recommended [4] as it gives the best estimation of the diameter of a 3-dimensional object (pore) from a 2-dimensional projection (image).

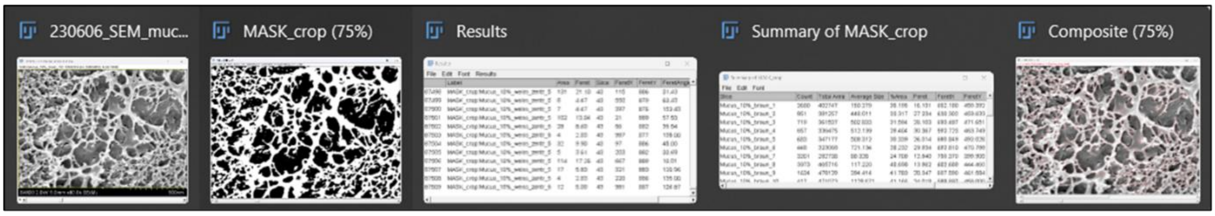

*Figure S1C: Output of the automated image analysis. From left to right: Original image (stack), binary result, Results table, Summary table, pore outline overlay for manual inspection.*

### Results

#### Additional cryo-SEM images

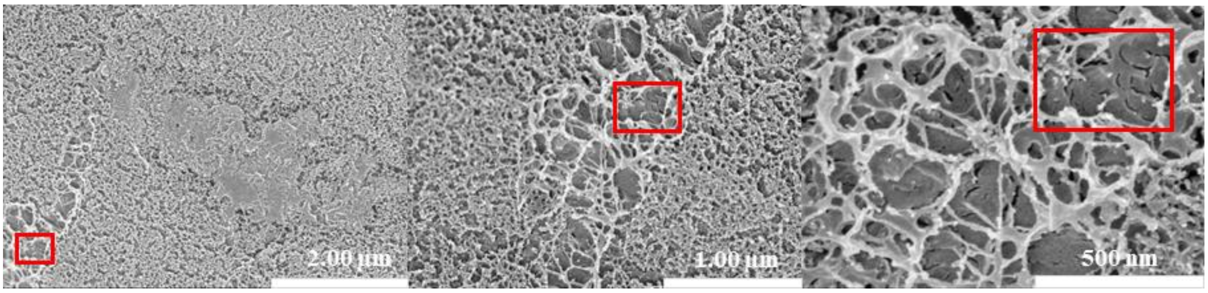


***Figure S2:*** *Cryogenic-scanning electron micrographs (cryo-SEM) at 20.000 ×, 40.000 × and 80.000 × magnification of a 100 mg/mL BSM solution, Batch-Nr. LOT 3776068. Higher resolution structures are outlined in red.*

As a supplement to Figure 2 A and B, additional SEM images are shown here, also of a 100 mg/mL BSM solution of batch no. LOT 3829388. The lowest magnification showed mainly fine-meshed areas, as can also be seen in Fig 2A with batch no. LOT 3776068. Of course, there werw also the larger meshed areas, as circled in red in the three ascending magnifications. Pore sizes of up to 500 nm are visible in this area. The violin plots (Fig 2C) clearly show the range of fine- and larger-mesh areas; the proportional distribution of small and large pores is also apparent.

#### Determination of the linear viscoelastic (LVE) region and the critical deformation

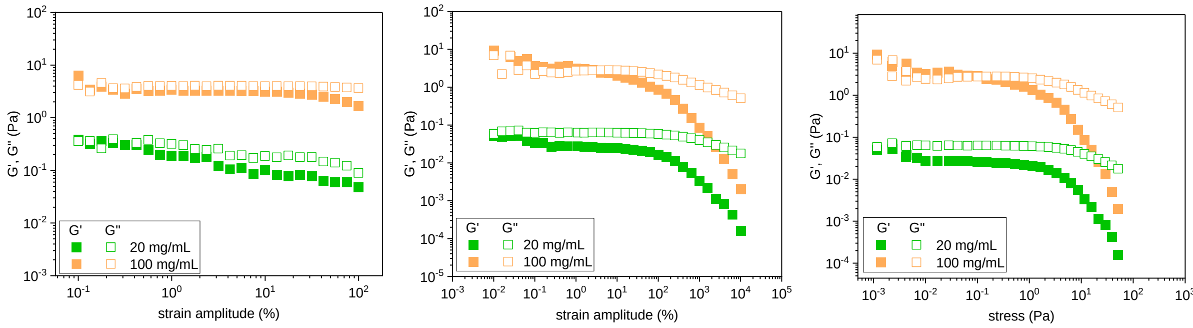


***Figure S3:*** *Storage modulus G' and loss modulus G'' as function of strain amplitude measured at 25 °C for BSM solutions of 20 and 100 mg/mL (frequency: 1 Hz); with a strain*

*amplitude from 0.1 to 100 % (left), 0.01 to 10000% (middle) and plotted against the stress (Pa) (right).*

The amplitude sweeps were performed at a constant frequency of 1 Hz. With an increase of the strain amplitude up to 100 % (left) the values for G’ and G’’ of the 100 mg/mL BSM solution remained at constant around 4 Pa for the loss modulus that is dominating over the storage modulus (G’ around 3 Pa). The situation with a 20 mg/mL BSM solution is somewhat different, because here the values for G' and G" did not behave so constantly over the measured amplitude range but decreased slightly with increasing amplitude. However, G" remained the dominant modulus (G'=0.15 Pa and G"=0.3 Pa at 0.1 % strain). The values of the moduli were less stable overall and have more dispersion compared to the 100 mg/mL solution, which was due to the lower concentration of the BSM solution. Based on these measurement results, the linear viscoelastic range was determined and the amplitude, which was then used for the subsequent frequency sweeps, was set to 1 %.

Furthermore, additional amplitude sweeps were performed with 20 mg/mL and 100 mg/mL BSM solution, where the amplitude was increased to higher values up to 10000% (middle). These showed a decrease for the storage modulus G' from an amplitude of 100 %. This value represents the critical deformation at which the mucus network is irreversibly changed. The corresponding yield stress value was around 0.7 Pa (right). The decrease of the loss modulus occurs with a lower slope and can be observed at a deformation between 100 and 1000 % while G" remains dominant over G’ (middle).

#### Concentration Dependence of the Viscoelastic properties of BSM solutions

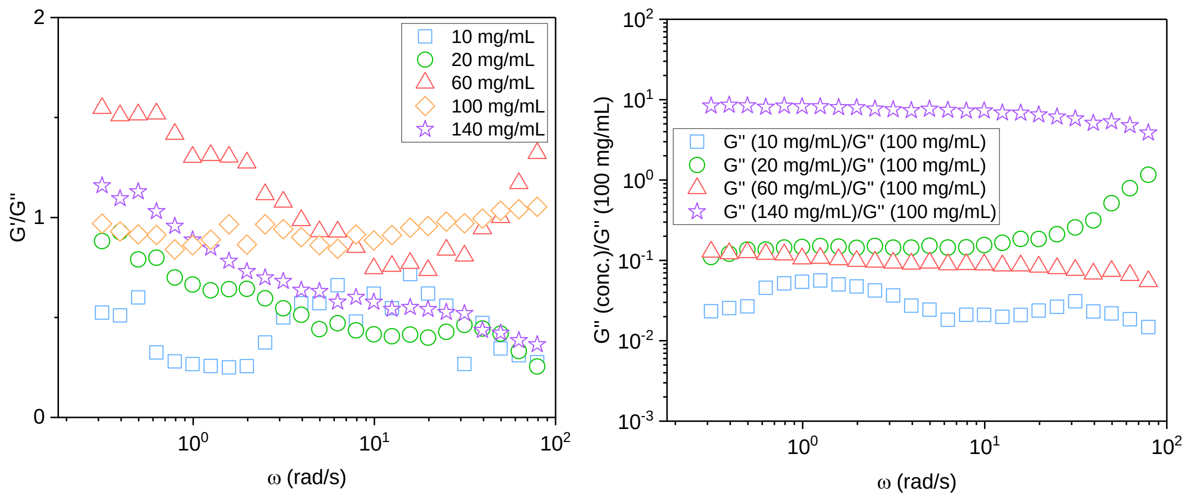

***Figure S4:*** *Ratio of G'/G'' of each concentration over the angular frequency (left); the loss modulus of each concentration is divided by the loss modulus of the 100 mg/mL concentration (right).*

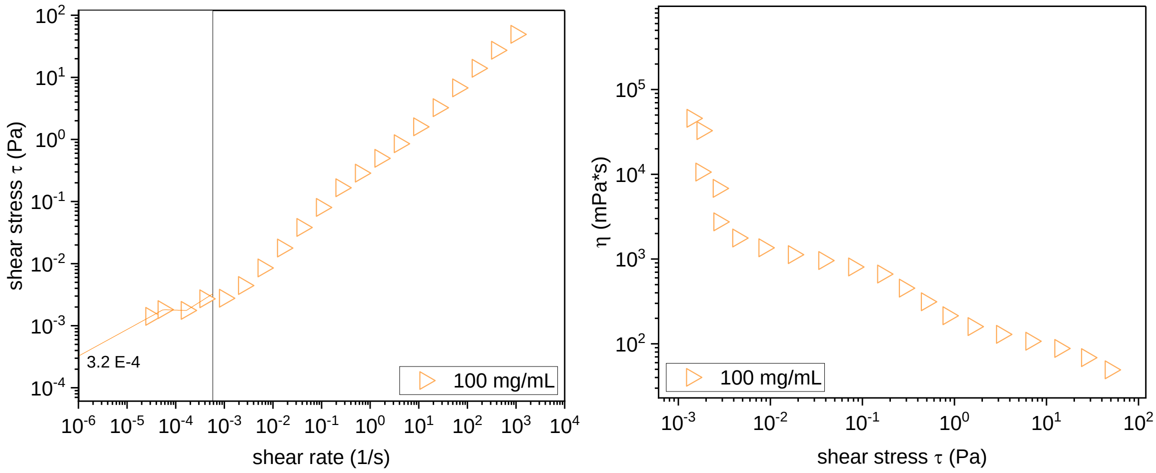


***Figure S5:*** *Shear stress τ (Pa) as a function of shear rate (1/s) (left) as well as shear viscosity η (mPa*s) as function of shear stress (Pa) (right) for a 100 mg/mL BSM solution. Data shown are the mean of n=3 measurements at this concentration.*

When performing the viscosity measurement at low shear rates (0.00001 1/s), an increase in the viscosity of a 20 mg/mL and 100 mg/mL BSM solution was observed. At the lowest shear rate of 0.00001 1/s, the viscosity of the 20 mg/mL BSM solution was 20000 mPa*s (20 Pa*s) and the 100 mg/mL BSM solution is 200000 mPa*s (200 Pa*s). A zero-shear viscosity could not be determined for this system.

The decrease in viscosity was more pronounced than for the measurements shown in Figure 3B, where measurements were taken at less low shear rates. A measurement from low to high shear rates and then back again showed a concordance of viscosity values and no hysteresis-like progression.

Additionally, the shear stress (Pa) was plotted as a function of the shear rate (1/s) (middle). By extrapolating the measured data from the yield curve to the point where it intersects with the shear stress axis, the yield point can be approximately determined, with a shear stress value of 3.2 $x10^{-4}$ Pa.

#### Effect of additives on the Viscoelastic properties of BSM solutions

##### Sodium chloride

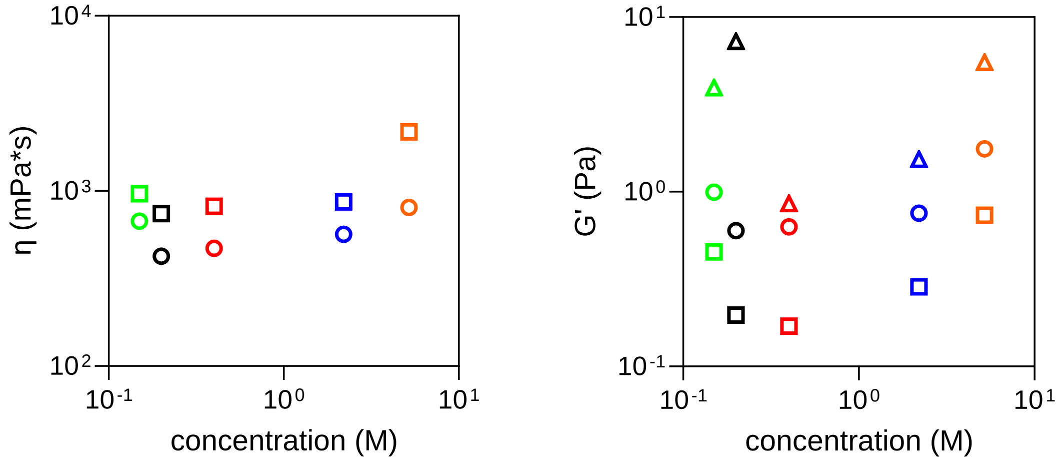


***Figure S6:*** *Viscosity η at two representative shear rates (0.1 1/s [squares] and 1.0 1/s [circles]) for each NaCl concentration (left). The values for the storage modulus for each NaCl concentration are shown at three different frequencies (0.1 Hz [squares], 1.0 Hz [circles] and 10.0 Hz [triangles]) (right).*

#### Calcium chloride

A

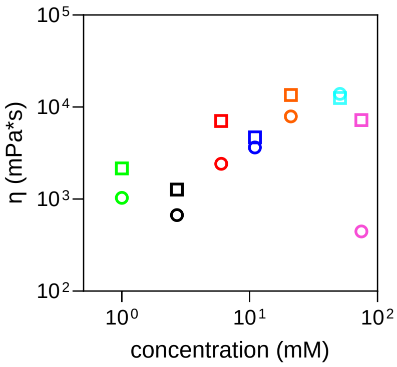


B

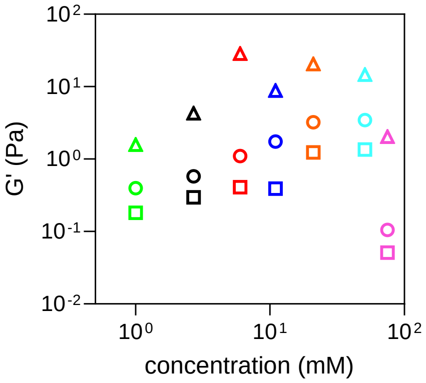


C

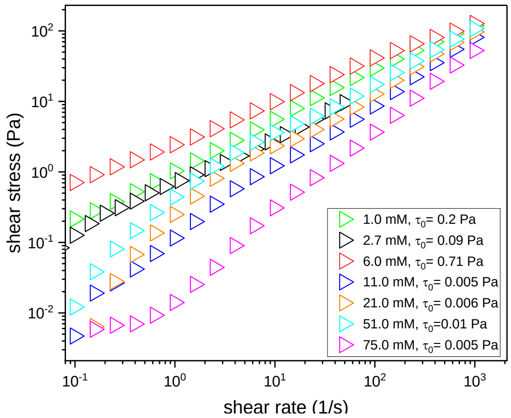


***Figure S7:*** *Viscosity $\eta$ at two representative shear rates (0.1 1/s [squares] and 1.0 1/s [circles]) for each $CaCl_2$ concentration (A). The values for the storage modulus for each $CaCl_2$ concentration are shown at three different frequencies (0.1 Hz [squares], 1.0 Hz [circles] and 10.0 Hz [triangles]) (B). Shear stress $\tau$ (Pa) as a function of shear rate (1/s) and the yield point $\tau_0$ of each $CaCl_2$ concentration (C).*

#### Lysozyme

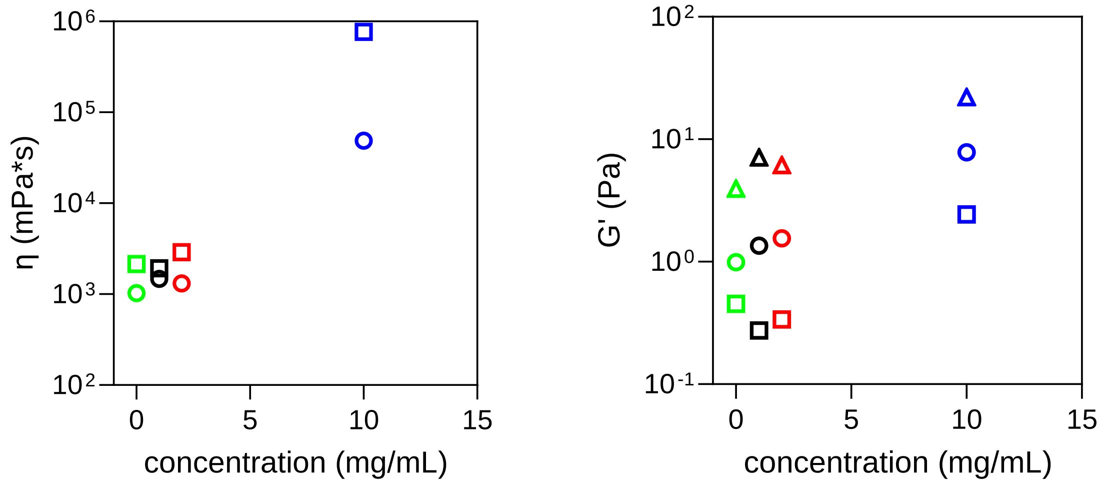


***Figure S8:*** *Viscosity η at two representative shear rates (0.1 1/s [squares] and 1.0 1/s [circles]) for each Lysozyme concentration (left). The values for the storage modulus for each Lysozyme concentration are shown at three different frequencies 0.1 Hz [squares], 1.0 Hz [circles] and 10.0 Hz [triangles]) (right).*

#### DNA

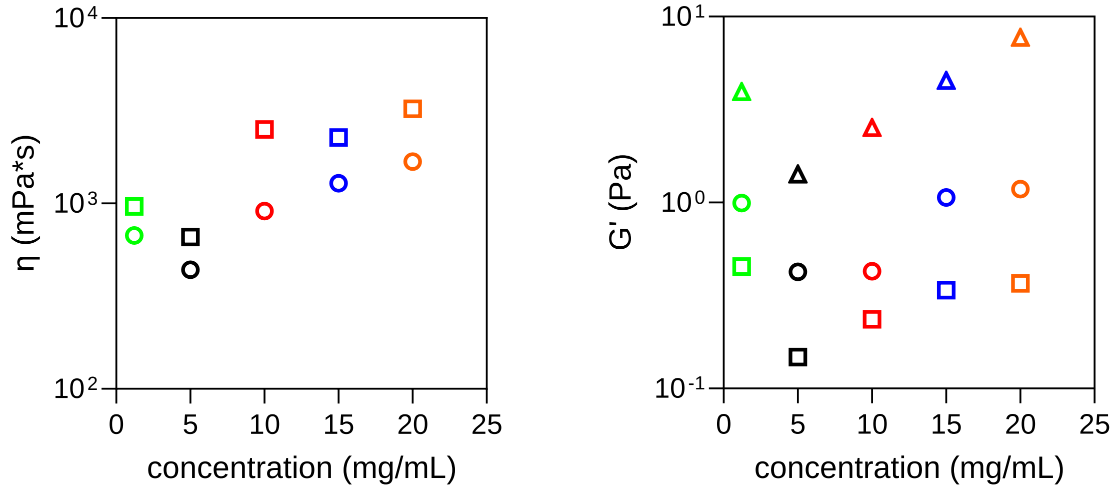


***Figure S9:*** *Viscosity η at two representative shear rates (0.1 1/s [squares] and 1.0 1/s [circles]) for each DNA concentration (left). The values for the storage modulus for each DNA concentration are shown at three different frequencies (0.1 Hz [squares], 1.0 Hz [circles] and 10.0 Hz [triangles]) (right).*

#### Detailed analysis of the rheological data

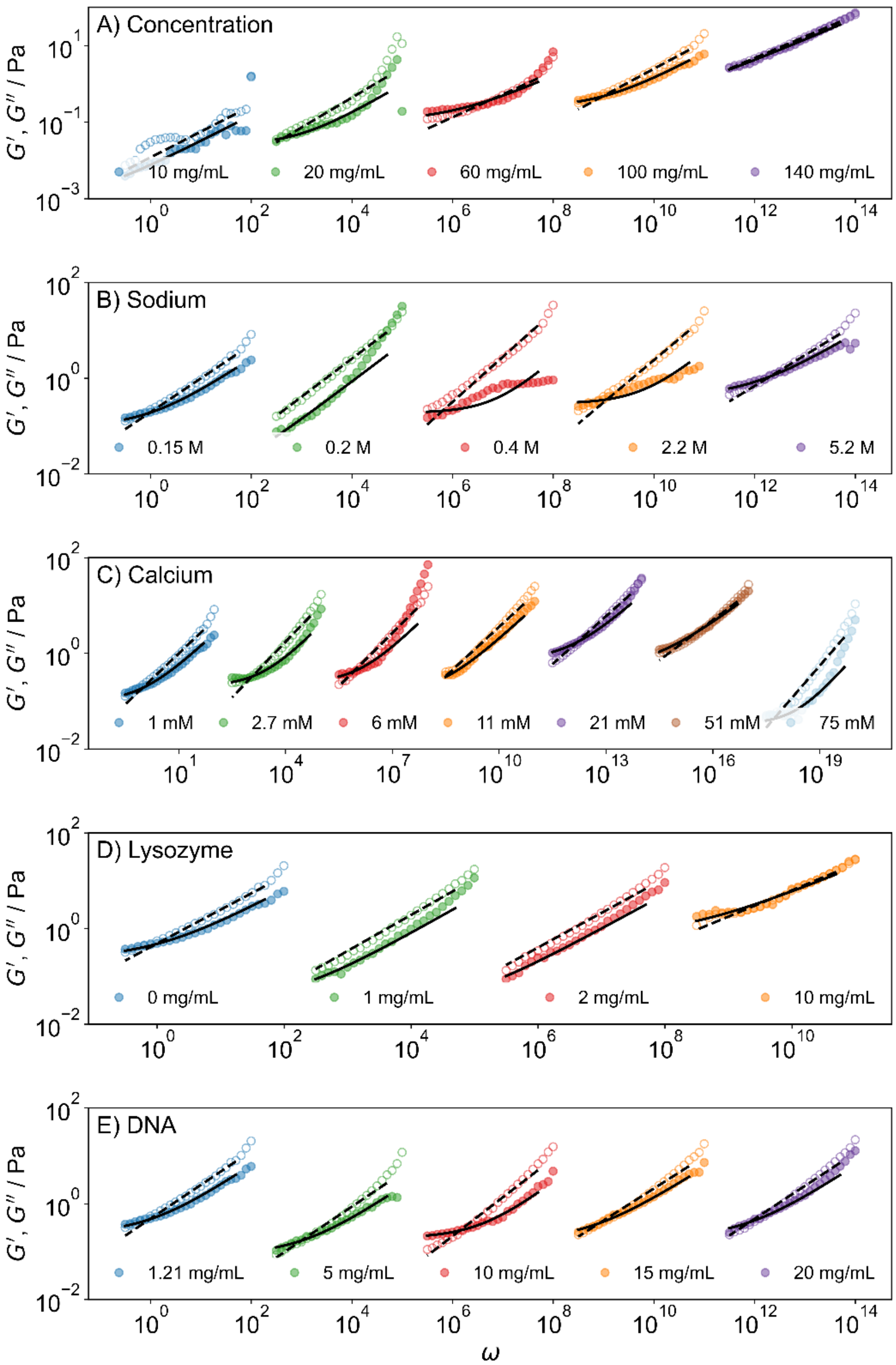


***Figure S10:*** *Storage modulus G' and loss modulus G'' as function of angular frequency (shifted by $10^3$ rad/s relative to each other); experimental data from rheological measurements (circles) fitted with the Fractional Kelvin-Voigt Model (black lines) for A) the five different concentrated BSM solutions, B) the BSM solutions with five different concentrations of added sodium chloride, C) the BSM solutions with seven different concentrations of added calcium chloride, D) the BSM solutions with five different*

*concentrations of added DNA, E) the BSM solutions with four different concentrations of added Lysozyme. The angular frequency of the first sample in each series is in rad/s.*

When examining the various concentrations of BSM solutions (Figure S10A), an increase in the viscoelastic moduli with increasing frequency can be seen. However, data obtained for low concentrated solutions (10 and 20 mg/mL) displayed a scattering pattern and exhibited a strong increase in the moduli at high frequencies, while the higher concentrated solutions showed a lower slope of the moduli with increasing frequency. The FKVM model with constant $\beta = 0$ demonstrated a good fit for the data (solid line = G' and dotted line = G"), describing the frequency-dependent viscoelastic behavior effectively, particularly at higher BSM concentrations. The fit works well, even when $\beta$ is kept constant at 0, making the interpretation of the model parameters easier. As a comparison, the fits with varying $\beta$ are shown in Figure S11.

The rheological data for BSM solutions with added sodium chloride (Figure S10B) fitted with the FKVM showed a small to negligible difference in viscoelastic moduli with increasing sodium concentration. Overall, the FKVM fitted values offered a close description of the behavior of the sodium BSM solutions.

With increasing concentration of added calcium chloride to the BSM solution (Figure S10C) we observed slightly increasing values for the viscoelastic moduli, except for the highest added calcium chloride concentration (75 mM). The FKVM models provided good results to describe the experimentally determined data and reflected well the crossover points of G' and G" observed at some concentrations.

As the concentration of lysozyme in a BSM solution (Figure S10D) increased, there seemed to be minimal alteration in the viscoelastic moduli of the system, except for the highest concentration of 10 mg/mL lysozyme. At this level, there is an increase in moduli and nearly equal distribution of the elastic and viscous fractions in the system. This outcome can be accurately represented by the FKVM.

The addition of DNA to a BSM solution (Figure S10E), even a rather high concentration of 20 mg/mL DNA, did not seem to have any effect on the viscoelastic properties of BSM. This is not so surprising as one may at least not expect any significant electrostatic interactions between the negatively charged BSM and the negatively charged DNA. The FKVM fitted data align well with the experimental data.

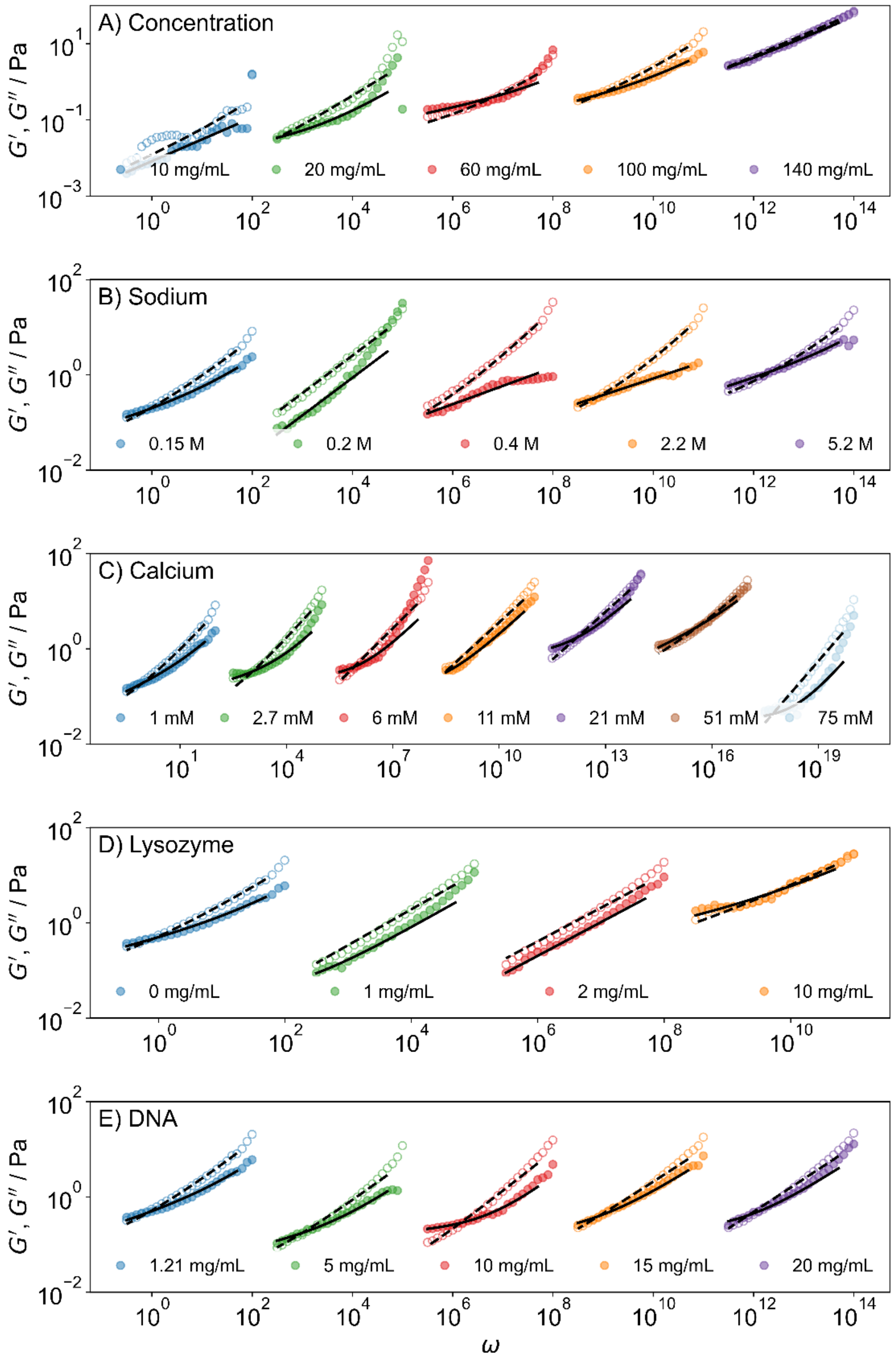


***Figure S11:*** *G' (filled circles) and G'' (empty circles) data from rheological measurements fitted with the FKVM (black lines) with varying β for BSM solutions with varying concentration of A) BSM, B) sodium, C) calcium, D) lysozyme and E) DNA. The angular frequency of the first sample in each series is in rad/s. Subsequent samples are shifted by* $10^3$ *rad/s to ensure readability.*
